# Bioinspired Oxidized mRNA Lipid Nanoparticles for *Ex Vivo* Engineering of Chimeric Antigen Receptor Macrophages Targeting Solid Tumors

**DOI:** 10.1101/2025.06.07.658363

**Authors:** Alvin J. Mukalel, Tina Tylek, Erin O’Brien, Caitlin Frazee, Hannah C. Geisler, Jacqueline Li, Ajay S. Thatte, Hannah C. Safford, Christian G. Figueroa-Espada, Mohamad-Gabriel Alameh, Alex G. Hamilton, David Mai, Neil C. Sheppard, Carl H. June, Drew Weissman, Kara Spiller, Michael J. Mitchell

## Abstract

Solid tumors remain difficult to treat via conventional and novel therapeutic strategies. Immunotherapies such as chimeric antigen receptor T (CAR-T) cell therapy have been remarkably effective in treating hematological cancers, but their efficacy is limited in solid tumors. Recently, CAR macrophages (CAR-Ms) have emerged as a promising solid tumor immunotherapy, primarily for their intrinsic tumor infiltration and effector functions. However, CAR-Ms are engineered using viral transduction, which is associated with aberrant immunogenicity and toxicity. To overcome these challenges, we developed a bioinspired oxidized lipid nanoparticle (LNP) platform for mRNA-based engineering of human CAR-Ms. A library of 24 ionizable lipids was synthesized, formulated into LNPs, and screened for delivery to human macrophages. The composition of the top LNP was subsequently optimized using an orthogonal design of experiments (DoE) and physicochemical properties, such as size and mRNA encapsulation, were tuned via optimization of microfluidic mixing parameters, yielding a particle that significantly outperformed a gold standard C12-200 LNP. Utilizing small molecule and antibody inhibitors, we demonstrate that uptake of optimized LNPs into macrophages is driven by apolipoprotein E (ApoE) independent macropinocytosis, which is further supported by potent extrahepatic spleen tropism upon intravenous administration to mice. Lastly, we demonstrate the translatability of this LNP platform and utilize it to engineer functional primary human HER2-CAR-Ms *ex vivo* with potent antigen-specific tumor killing, validated in an *ex vivo* co-culture with ovarian cancer cells. This bioinspired oxidized LNP platform can potentially be utilized to engineer a range of human CAR-M immunotherapies to treat various types of solid tumors.

## Introduction

Solid tumors are a longstanding challenge to treat using conventional and immunotherapies.^1,2^ Chimeric antigen receptor (CAR) T cell immunotherapy, in which patient T cells are isolated, virally engineered to express a tumor-targeted CAR, and reinfused back into patients, has demonstrated profound efficacy in treating hematological malignancies. However, solid tumors present various physical, chemical, and signaling barriers that have hindered the efficacy of CAR T cells.^3,4^ Recent preclinical strategies to enhance CAR T cell efficacy against tumors have introduced additional therapeutic modalities such as vaccines and oncolytic viruses to enhance T cell infiltration and crosstalk with the innate immune system.^5,6^ However, these approaches increase the complexity of the therapeutic which in turn prolongs the regulatory process for clinical translation. Similarly, immune checkpoint blockade (ICB) has dramatically improved survival rates in treating solid tumors. However, its use is restricted to only certain patients by factors such as the heterogenous expression of targetable molecules such as PD-1 and CTLA-4, which vary across patients and must be expressed at high enough levels to be therapeutically relevant. Even if these criteria are met, the response rates can vary, and it is estimated that only 49% of patients are eligible for checkpoint inhibitor blockade, and only 12% of those might respond.^7^ Thus, despite the objective successes of the aforementioned immunotherapies, newer modalities are required to treat solid tumors.

Macrophages, a type of phagocytic immune cell, have long been considered a promising therapeutic target for treating solid tumors.^8–12^ In addition to being key regulators of the innate immune system, macrophages have an intrinsic ability to infiltrate solid tumors and, in certain tumor types, can comprise up to 50% of the tumor mass.^13^ Once in the tumor, macrophages typically adopt immunosuppressive phenotypes that support tumor growth and metastasis through the surface expression of checkpoint inhibitors (*e.g.,* PD-1, PD-L1, PD-L2, and CTLA-4 ligands)^11,14,15^ and the secretion of proangiogenic factors (*e.g.,* VEGF, PDGF, Ang1, and TIE2)^16–18^ and immunosuppressive cytokines (*e.g.*, IL-4, IL-10, and TGFβ).^19–21^ Thus, prior macrophage-targeted therapies have aimed to block these behaviors using targeted antibodies,^22–26^ induce repolarization towards immunostimulatory phenotypes,^27,28^ and reduce expression of checkpoint molecules.^11^ Macrophages bridge the innate and adaptive arms of the immune system by performing functions in addition to phagocytosis including antigen presentation, T cell stimulation, and cytokine secretion: processes essential to generating tumor immunity.^9,29^ Therefore, macrophages have emerged as an immunoengineering target where, given the proper stimuli, their tumor infiltrating and immunoregulatory properties can be leveraged for novel solid tumor immunotherapies.^30–35^

One such macrophage-targeted therapy is CAR-macrophage (CAR-M) therapy, which has demonstrated significant promise as a new solid tumor immunotherapy.^35^ Engineered macrophages can directly participate in CAR-directed tumor cell killing via phagocytosis and further engage the immune system through antigen presentation and potent cytokine production. Altogether, CAR-Ms are capable of orchestrating a broad and multifaceted anti-tumor immune cascade.^10,35^ Because of their potential to treat solid tumors, HER2-targeted CAR macrophages (HER2-CARMs) were granted a fast-track designation by the FDA in 2021, and are currently being studied in phase 1 clinical trials for HER2-overexpressing solid tumors (NCT04660929).^36^

Although CAR-Ms have rapidly transitioned from preclinical research to clinical trials, they are still encumbered by engineering limitations.^37^ *Ex vivo* macrophage engineering has been a significant challenge, as they are terminally differentiated, non-dividing cells that specialize in the degradation of foreign extracellular materials and pathogens; thus, exogenous gene transfer via viral and non-viral vectors is difficult and requires carefully designed platforms.^38–42^ Currently, CAR-Ms are engineered to express CAR using lentiviruses or adenoviruses (AdVs), which utilize cognate receptor-ligand interactions to trigger endocytosis and subsequently induce long-term and stable expression of the therapeutic CAR cassette.^35,42^ However, viral-induced CAR expression, especially in the context of solid tumor immunotherapy, is strongly associated with significant adverse effects, and even death.^43–45^ These adverse effects are often due to the scarcity of tumor-specific antigens (TSAs) that could provide a means for exclusive targeting of the cancerous tissue. Instead, tumors are targeted using common antigens that are not tumor-specific and rather overexpressed in cancerous tissue relative to the rest of the body, which can lead to on-target, off-tumor toxicity.^46–48^ Further, viral vectors can activate intracellular inflammatory pathways in engineered macrophages and induce a highly durable inflammatory phenotype due to their relatively prolonged residence within the cell. While this may lead to desirable immunostimulatory effects in the short term, chronic inflammatory signaling via molecules such as TNF-α and IL-6 can lead to an accrual of DNA damage, differentiation of suppressive immune cells, and promote therapeutic resistance, together leading to cancer growth and metastasis.^49–55^ Thus, the viral vector itself and expression of viral therapeutic cassettes are associated with several adverse effects.

As a non-viral alternative, mRNA therapeutics are rapidly garnering interest due to their potency, non-integrating nature, and potential for tunability. As opposed to viruses, mRNA can be synthesized at scale without cell cultures, does not interact with the host genome, and is only transiently expressed.^56–60^ For this reason, mRNA has found a particular niche in CAR-based therapies, as it enables precise and well-defined expression kinetics that can obviate the on-target, off-tumor toxicity associated with long-term viral CAR engineering.^61–65^ However, mRNA is a large, anionic macromolecule that cannot readily cross cell membranes on its own. Current clinical standards utilize electroporation (EP) for *ex vivo* engineering, where the cell is transiently permeabilized using electrical pulses, allowing mRNA to enter the cell. However, this technique can lead to cell toxicity and undesirable phenotypic changes, and EP itself is severely limited in terms of its scalability and *in vivo* administration.^66,67^ Thus, for potent mRNA delivery, the cargo must be encapsulated in a delivery vehicle to simultaneously protect it from enzymatic degradation and to facilitate transport across the cell membrane to the cytosol where it mediates its therapeutic function.^68^

Lipid nanoparticles (LNPs) are the most clinically advanced mRNA delivery vehicle, as the Onpattro siRNA-LNP therapeutic developed by Alnylam has been FDA-approved since 2018 and the two COVID-19 mRNA LNP vaccines developed by Pfizer/BioNTech and Moderna received full FDA approval in 2022. LNPs are composed of a pH-responsive ionizable lipid that becomes positively charged at acidic pH, a membrane-fusogenic phospholipid, cholesterol, and a polyethylene glycol-lipid (PEG-lipid) conjugate that promotes stability in the biological environment.^68–73^ Combining these lipids allows for efficient encapsulation of the mRNA cargo, homing of the mRNA-LNP to the target tissue, and intracellular release of the mRNA in the desired cell type. Together, LNPs are well suited to overcome the barriers to intracellular mRNA delivery and offer a promising platform for generating CAR-M therapies.^74–76^

Here, we designed an mRNA delivery platform for human macrophages through three rounds of LNP optimization: (1) the ionizable lipid structure, (2) the LNP excipient formulation ratios, and (3) the LNP physicochemical properties (**Figure 1**). A library of 24 ionizable lipids was synthesized, formulated into mRNA-containing LNPs, and screened for delivery of luciferase-encoding mRNA to PMA-differentiated human THP-1 macrophages. Through the lipid screen, we identified a top LNP, the bioinspired C16-C lipid, with high potency and minimal toxicity. The excipient ratios of the C16-C formulation were then optimized using an orthogonal design of experiments (DoE) approach, yielding an optimized B15 formulation with 13-fold higher delivery compared to the initial C16-C formulation. Next, we assessed a range of microfluidic mixing conditions, enabling us to further tailor the hydrodynamic radius and mRNA encapsulation of the LNP formulation to enhance mRNA delivery to macrophages. Using the fully optimized mRNA-LNP we studied the mechanisms by which LNP uptake occurs — observing that uptake is largely driven by ApoE-independent macropinocytosis — and the effect of LNP treatment on macrophage phenotype, finding that the LNPs themselves generate a mild, but insignificant increase in proinflammatory gene expression. Lastly, to emphasize the potential for translation with this optimized LNP platform, we demonstrated potent *ex vivo* mRNA transfection in both granulocyte macrophage colony stimulating factor (GMCSF) and macrophage colony stimulating factor (MCSF) models of primary human macrophages, and further engineered functional primary human HER2-CAR macrophages which were evaluated using an *ex vivo* tumor co-culture killing assay.

**Figure 1.**
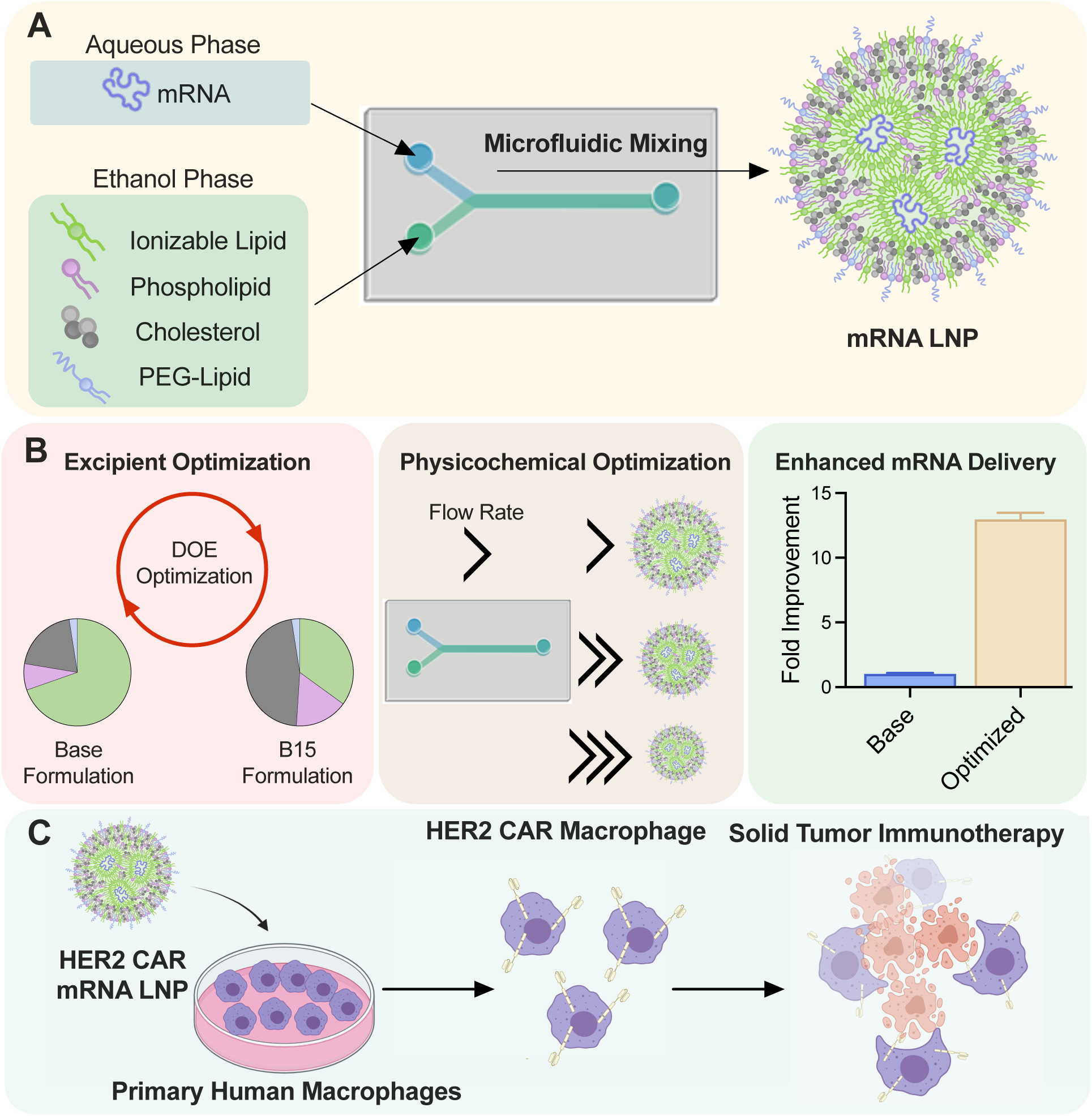
Project Overview. (A) An ionizable lipid library containing bioinspired oxidized ionizable lipids was formulated into ionizable lipid nanoparticles (LNPs) via microfluidic mixing of a lipid-containing ethanol phase with an mRNA-containing aqueous phase and screened for delivery to THP-1 macrophages. (B) The excipients and physicochemical characteristics of the top LNP formulation, an oxidized LNP, were subsequently optimized using a design of experiments (DoE) and microfluidics to yield an optimized formulation 13-fold more potent than the base formulation. (C) The optimized LNP formulation was used to engineer functional HER2-targeted CAR macrophages *ex vivo*, capable of killing HER2+ ovarian cancer cells and demonstrating the LNPs platform’s potential as a solid tumor immunotherapy.

## Results and Discussion

### Screen of piperazine-derived ionizable lipids in THP-1 macrophages

In this study, a library of 24 ionizable lipids were synthesized based on a gold standard parent molecule, C12-200 (**Figure 2A-C**).^77^ A subsection of the library was designed to include varying degrees of oxidation into the parent 200 polyamine core molecule, taking inspiration from macrophages’ intrinsic affinity for the uptake of naturally occurring oxidized lipid structures.^78,79^ These bioinspired polyamine cores included cores C, F and G (**Figure 2B**). Ionizable lipids were synthesized using one-pot S_N_2 chemistry in which polyamine cores were reacted with epoxide-terminated 12, 14, or 16 alkyl chains (C12, C14, C16). Synthesized ionizable lipids were then combined with cholesterol, DOPE, and C14-PEG 2000 (PEG-Lipid) in a single ethanol phase at molar ratios of 35:46.5:16:2.5 and mixed with an mRNA-containing aqueous phase (10 mM citrate buffer, pH = 4) in a microfluidic mixing device.^80^ Dynamic light scattering (DLS) measurements demonstrated that all LNPs in the library had intensity-based diameter between 60 and 100 nm with polydispersity index (PDI) below 0.3, indicating a monodispersed mixture, with all mRNA concentrations in LNP solutions >30 ng/μL (**Supplementary Table 1**). LNPs encapsulating luciferase-encoding mRNA were evaluated for potency in human THP-1 monocytes, a common model for peripheral blood mononuclear cell (PBMC) derived monocytes and macrophages, that were differentiated into macrophages by a 48 h incubation in full growth media supplemented with phorbol 12-myristate 13-acetate (PMA, 10 ng/μL).^81^ This *in vitro* screen identified 4 LNPs with more potent mRNA delivery to macrophages compared to the gold standard parent ionizable lipid, C12-200 (**Figure 2D**). We identified a trend that ionizable lipids with longer tails (C14 and C16) and internal sites of oxidation between amines (cores C, F, G) tended to have greater mRNA transfection in THP-1 macrophages (**Figure 2D**). Using a relative hit rate analysis, a method previously employed to derive ionizable lipid structure-activity relationships, we identified that bioinspired oxidized ionizable lipids were overrepresented among the hits from this initial screen, and the top LNPs all contained bioinspired ionizable lipids, demonstrating the utility of this design approach for engineering LNPs for macrophage delivery (**Figure 2E**).^82^ Other ionizable lipid structures screened in the library with various structural features such as internal branching and increased number of ring structures, piperazine groups, or amines enabled minimal mRNA delivery to THP-1s (**Figure 2E**). The four lead lipids (C14-C, C16-C, C14-F, C16-F) and one additional lipid (C16-G), were further studied to evaluate their dose-dependent delivery, where the C16-C lipid was found to be the most potent while showing minimal toxicity at a dose of 500 ng mRNA per 50k cells (**Figure 2F, Supplementary** Figure 1).

**Figure 2:**
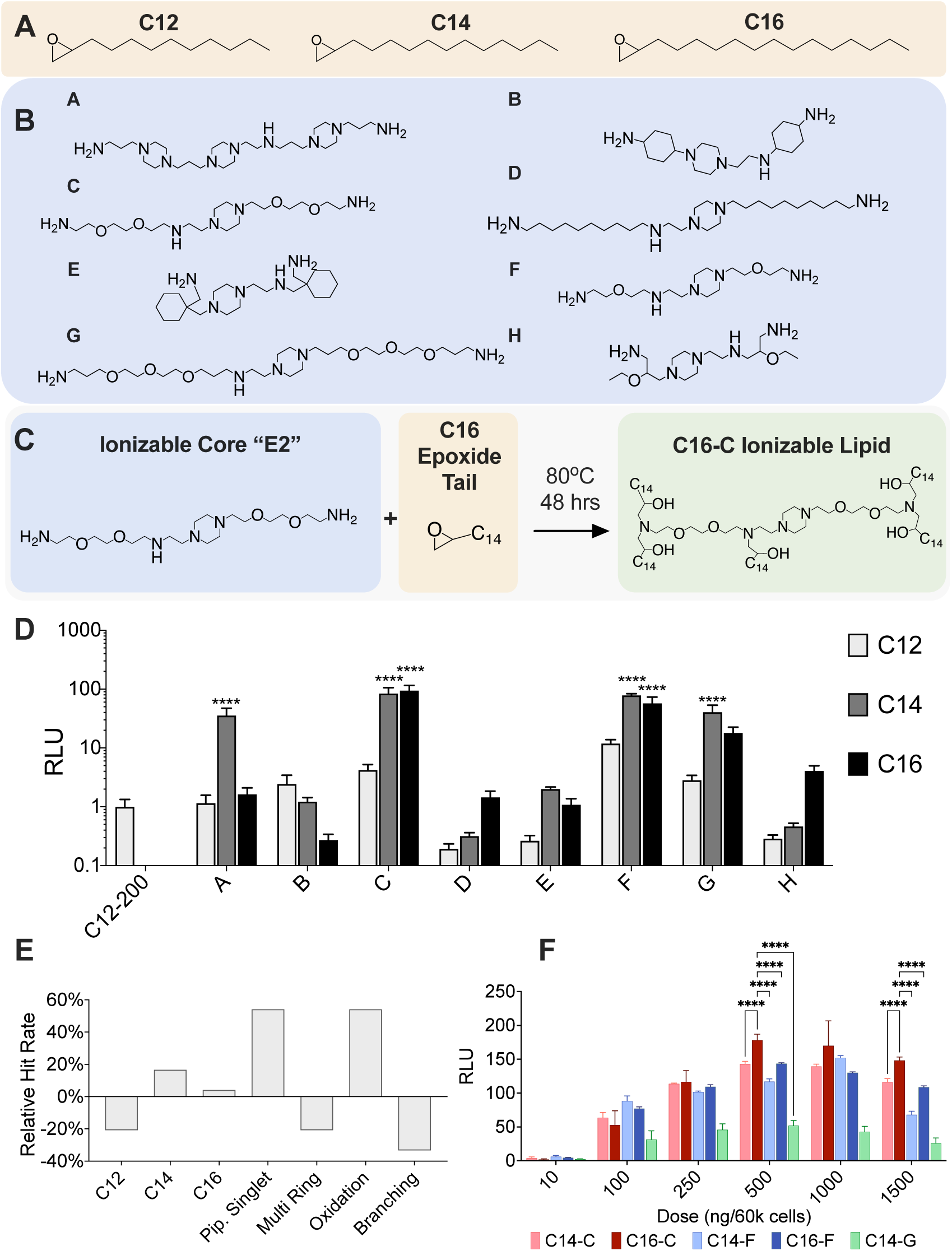
Lipid screen of 24 ionizable lipids identifies the bioinspired oxidized C16-C lipid capable of potent mRNA delivery to human THP-1 macrophages. (A) Epoxide-terminated tails and (B) polyamine cores were combined using an S_N_2 reaction to formulate 24 unique lipids. (C) Example S_N_2 reaction for the synthesis of C16-C ionizable lipid. (D) PMA-differentiated human THP-1 macrophages were treated with luciferase mRNA-LNPs at a dose of 250 ng/ 50k cells. Luminescence was measured 24 hours later, normalized to cells treated with a C12-200 gold standard and compared to the C12-200 group using a 1-way ANOVA with *n* = 5 biological replicates. (E) Relative hit rate analysis, where a hit is defined as RLU 5-fold higher than C12-200 of the ionizable lipid screen in (D), identifies bioinspired oxidized ionizable lipid design as a potential driver for LNP-mediated mRNA delivery to macrophages. (F) PMA-differentiated THP-1 macrophages were treated with the top five luciferase mRNA-LNPs identified in (D) at the indicated doses. Luminescence was measured 24 hours later, normalized to untreated cells and compared using a 2-way ANOVA with Holm-Sidak correction with *n* = 4 biological replicates. * p < .05, ** p < .01, *** p < .005, **** p < .001

### Optimization of C16-C LNP using orthogonal design of experiments (DoE)

After identifying the C16-C ionizable lipid as the lead candidate from the ionizable lipid library, we optimized the composition of the LNP using an orthogonal Design of Experiments (DoE).^83^ For four component systems such as these LNPs, varying the abundance of each component at 4 different levels would typically require 4^4^ = 256 independent formulations to be tested. However, an orthogonal DoE enables each parameter to be estimated independently of each other, allowing for the entire testable space of 256 formulations to be sampled with only 16 representative formulations (**Figure 3A**). The optimization was performed using a sequential two-step approach. In the first step, Library A, the base formulation was used as the center point and the additional formulations covered a wide range of excipient molar ratios around the base formulation to estimate which excipients had the most significant impact on mRNA delivery. Since macrophages are highly engaged in endocytosis and phagocytosis, we hypothesized that ionizable lipid content would be the strongest determinant of mRNA delivery, as the ability to escape from acidic subcellular compartments might be especially important for cytosolic delivery to this cell type. As such, Library A was designed to have a wide range of ionizable lipid content (5-50 molar ratio). The second library, Library B, was informed by trends elucidated by the screen of Library A, but at a narrower range of excipient molar ratios, thereby providing higher resolution in screening to identify an optimized formulation.

**Figure 3:**
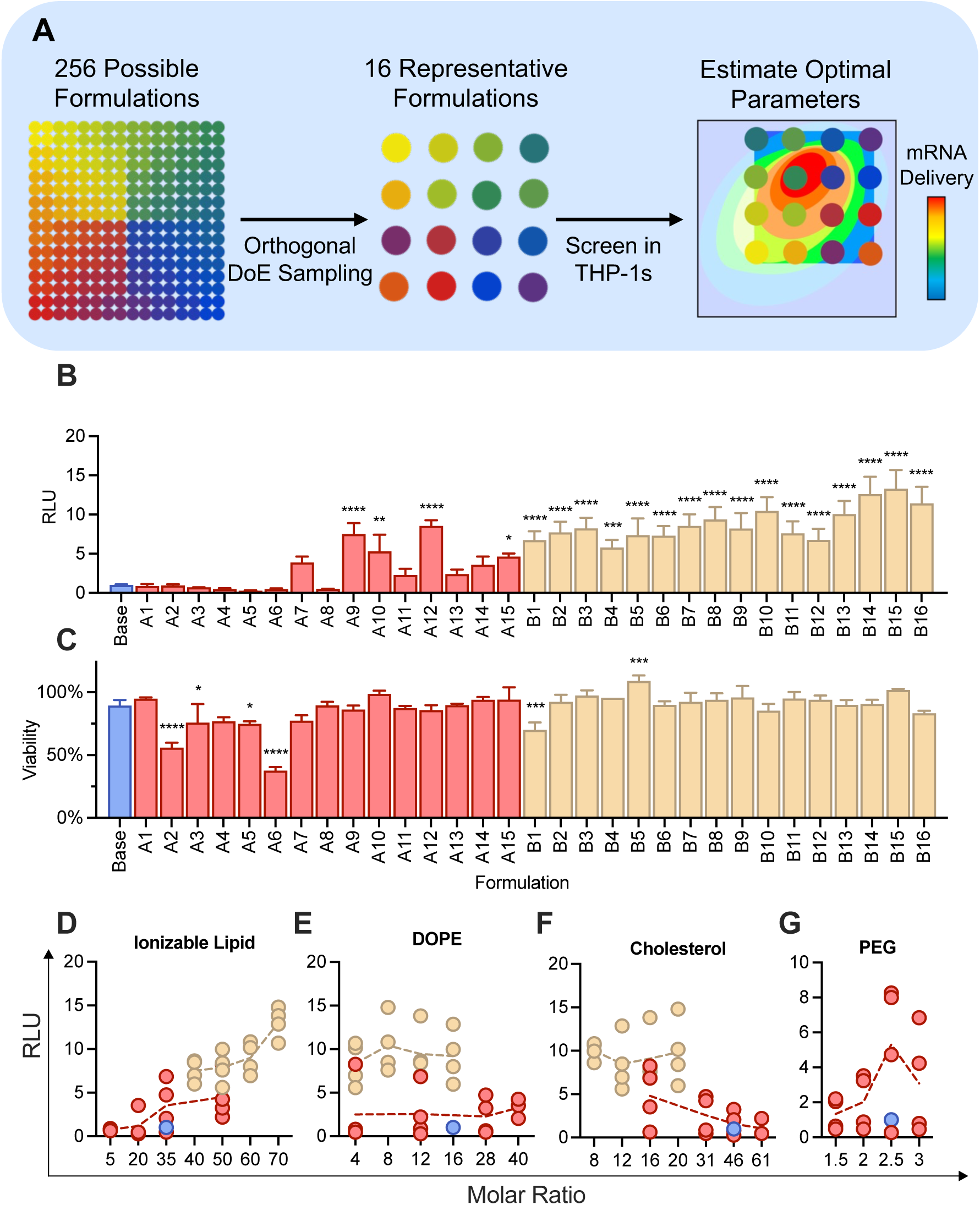
An orthogonal DoE identifies the B15 excipient formulation that enhances mRNA delivery to THP-1 macrophages 13-fold. (A) Sequential rounds of orthogonal DoE screening enabled the entire 256-formulation space to be represented as 16 unique formulations that can be rapidly screened *in vitro.* (B) PMA-differentiated THP-1 human macrophages were treated with the indicated formulation at a dose of 250 ng/50k cells. Luminescence was measured 24 hours later, normalized to the signal from the Base formulation, and analyzed using a one-way ANOVA with a Holm-Sidak correction with comparisons to the base formulation with *n* = 4 biological replicates. (C) PMA-differentiated THP-1 human macrophages were treated with the indicated formulation at a dose of 250 ng/50k cells. Viability was measured 24 hours later using a CellTiter Glo assay. Signal was normalized to untreated cells and analyzed using a one-way ANOVA with a Holm-Sidak correction with *n* = 4 biological replicates. (D-G) Scatterplots relating the abundance of each LNP component with mRNA delivery (RLU). Each formulation is represented as a circle, with the average represented as the dotted line. * p < .05, ** p < .01, *** p < .005, **** p < .001

Library A was designed with 16 formulations that varied in ionizable lipid content, DOPE, and cholesterol across a wide range of molar ratios; the formulated LNPs were evaluated for physical parameters such as size, PDI and mRNA concentration. 15 of the 16 LNP formulations in Library A met our criteria for desired physical parameters (mean size < 200 nm; PDI < 0.3; mRNA concentration > 10 ng/μL) (**Supplementary Table 4**); thus, these 15 LNPs were screened for mRNA delivery *in vitro*. For mRNA delivery and toxicity assays, THP-1s were treated with luciferase-mRNA LNPs at a dose of 250 ng mRNA per 50k cells. We identified four formulations, A9, A10, A12, and A15, with enhanced mRNA delivery to THP-1 cells, with the A12 formulation having an ∼8-fold increase in delivery compared to the base formulation, with no significant toxicity **(Figure 3B-C**). Interestingly, LNPs A7-A15, formulations with higher ionizable lipid content and lower phospholipid (DOPE) and cholesterol content, generally outperformed the base formulation. In comparison, LNPs A1-A6, formulations with lower ionizable lipid content, lower PEG-lipid content, and higher phospholipid (DOPE) and cholesterol content, demonstrated significant toxicity with equivalent or lower LNP potency compared to the base formulation.

We constructed our second DoE library, Library B, using the A12 LNP formulation as the center point, to explore a higher range of ionizable lipid content with lower molar ratios of phospholipid (DOPE) and cholesterol with a fixed PEG molar ratio (2.5 molar ratio). Library B greatly enriched mRNA delivery to THP-1 cells, with all 16 formulations significantly outperforming the base formulation and the highest performing formulation, B15, having a 13-fold increase in luminescence signal (**Figure 3B-C**). Interestingly, both rounds of DoE revealed positive trends between mRNA delivery and ionizable lipid content, and a negative correlation between mRNA delivery and cholesterol content, but no trend was found between mRNA delivery and phospholipid content (**Figure 3D-G**). These findings from the DoE screens support the hypothesis that ionizable lipid content is the major driver of intracellular delivery to macrophages *in vitro*. Although there were trends found in these studies, it is important to note the inherent limitations to these *in vitro* studies, as optimization for delivery *in vitro* and *in vivo* likely differ from each other. In this study, the top performing B15 formulation has low cholesterol content, whose primary role is to aid in LNP stability, which is more likely to be important under dynamic *in vivo* conditions.

### Optimization of the B15 LNP by carrier:cargo (wt:wt) formulation ratio

Although the excipient optimization of the B15 LNP yielded a high performing formulation with the C16-C ionizable lipid, we sought to determine whether optimizing formulation parameters, such as ionizable lipid:mRNA (wt/wt) ratio, could additionally impact LNP-mediated mRNA delivery to THP-1 macrophages. Recent studies have demonstrated a positive correlation between ionizable lipid:mRNA ratio (wt:wt) and mRNA delivery to macrophages *in vitro*, the result of which is more lipid per molecule of mRNA.^84^ We examined if this principle could be applied to the C16-C LNPs and if increasing the amount of the lipid in the formulation had any interaction with the composition of the LNP formulation (i.e. LNP excipient molar ratios).

Three formulations were selected from the previous LNP screens: the screening formulation (Base), the top formulation from Library A (A12), and the excipient-optimized formulation (B15) (**Figure 4A**). These were formulated at 6 ionizable lipid:mRNA ratios (wt/wt), increasing from the standard 10:1 (wt:wt) ratio to a maximum of 25:1 (wt:wt) while keeping the excipient ratios for each formulation constant. We observed improved delivery for the Base and A12 formulations at weight ratios > 10:1, with delivery reaching a maximum at higher weight ratios for the Base formulation (17.5:1) than the A12 formulation (15:1) (**Figure 4B-C**). In contrast, the B15 formulation showed the opposite trend. All B15 LNPs formulated at ratios greater than 10:1 had decreased luminescence, indicating decreased mRNA delivery (**Figure 4D**). Overall, formulations with lower ionizable lipid content had increased activity in macrophages at higher ionizable lipid:mRNA weight ratios. Conversely, formulations with higher ionizable lipid content had decreased delivery when formulated at higher ionizable lipid:mRNA weight ratios. From an excipient standpoint, increasing ionizable lipid content corresponds to proportionately lower DOPE, cholesterol, and PEG-lipid content. These results suggest a necessity of these excipients as the ionizable lipid:mRNA ratio increases, as they may help to stabilize the formation of mRNA-LNP and may help drive the formation of more LNPs that encapsulate mRNA. This is highlighted by the Base formulation, which has a higher abundance of excipients such as cholesterol, DOPE and PEG-lipid, having improved delivery at all increased weight ratios greater than 10:1. Although formulations B15 and A12 have similar amounts of C16-C ionizable lipid and DOPE, the A12 formulation has higher cholesterol and PEG-lipid content, potentially enhancing its stability and subsequently contributing to the observed increases in delivery at higher weight ratios not observed with the B15 LNP.

**Figure 4:**
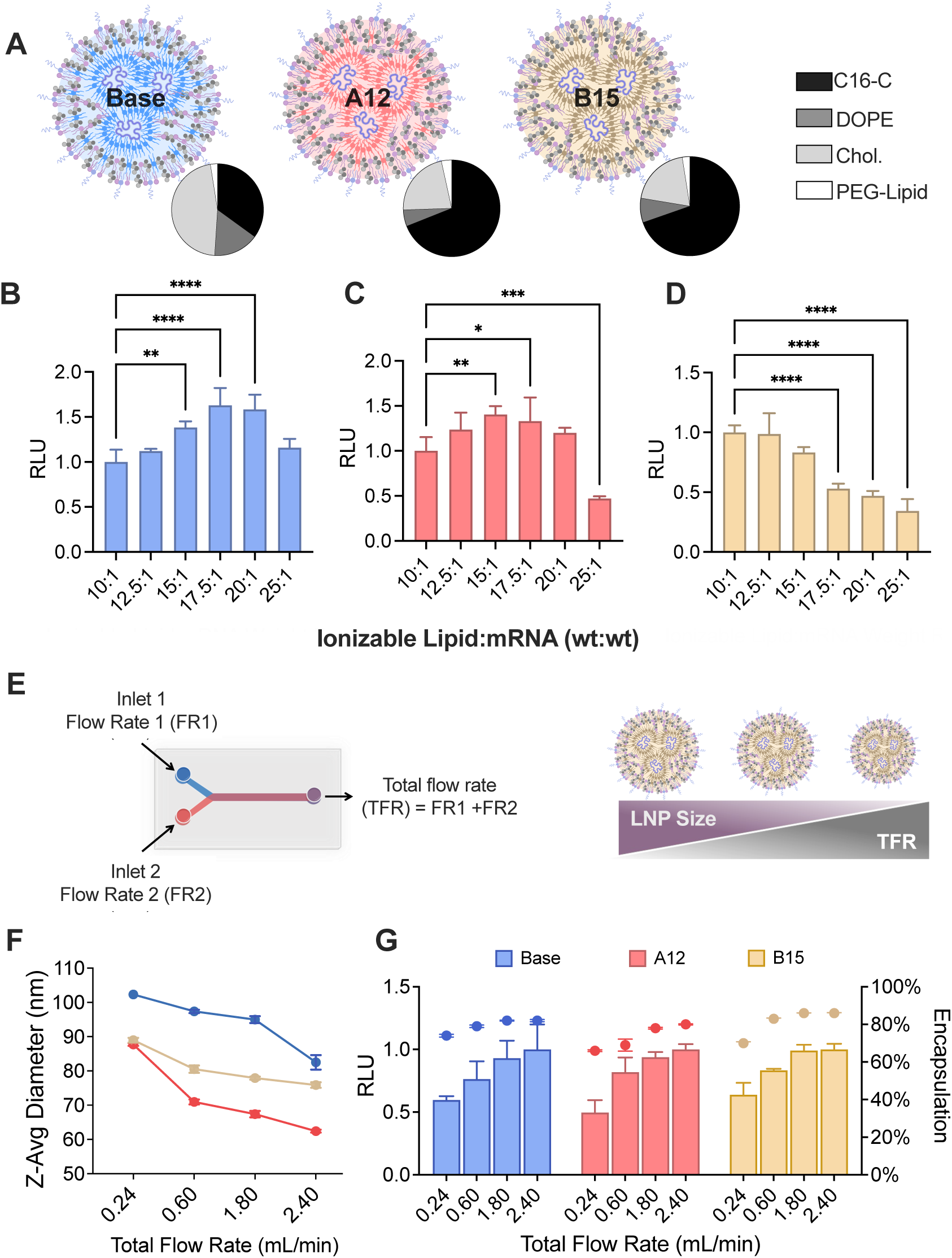
mRNA LNP delivery to macrophages can be tuned by adjusting LNP physicochemical parameters. (A) Excipient compositions (molar ratio) of the Base, A12, and B15 formulations selected for physicochemical optimization. (B-D) PMA-differentiated human macrophages were treated with the (B) Base, (C) A12, or (D) B15 LNPs formulated at varying ionizable lipid (C16-C):mRNA weight ratios at a dose of 250 ng/50k cells. Luminescence was measured 24 hours later, normalized to the 10:1 (wt:wt) group, and analyzed using an ordinary one-way ANOVA with a Holm-Sidak correction with *n* = 4 biological replicates. (E-F) LNP hydrodynamic radius has an inverse relationship with total flow rate through the microfluidic mixing device. Hydrodynamic radius (F) was measured using dynamic light scattering and represented as the average of 3 measurements. (G) mRNA Encapsulation efficiency (circles) was quantified using a Ribogreen Assay. PMA-differentiated human macrophages were treated with LNPs formulated at the indicated TFRs at a dose of 250 ng/ 50k cells for 24 hours. Luminescence was measured 24 hours later, normalized to the 2.4 mL/min TFR signal, and compared using a 2-way ANOVA with Holm-Sidak correction with *n* = 4 biological replicates. * p < .05, ** p < .01, *** p < .005, **** p < .001

### Microfluidic-based physicochemical optimization of the B15 LNP

Microscale mixing of fluids via microfluidic devices, such as the staggered herringbone mixers utilized here, enable precise control and formulation of LNPs.^80^ Previous studies have shown that total flow rate (TFR), where the TFR through the device is equal to the sum of the input flow rates, can be used to tune the size of LNPs, with lower TFRs yielding larger LNPs (**Figure 4E**).^80^ In the context of macrophage nanotherapies, size and morphology play a significant role in influencing particle intracellularization and the subsequent delivery of therapeutic cargos.^85–87^ Thus, we explored formulation-dependent relationships between TFR, LNP size, and mRNA delivery in THP-1 macrophages.

We first confirmed that varying TFR enabled us to tune the size of the LNPs independently of the excipient molar ratios (**Figure 4F**). It was also noted that decreasing TFR appeared to have a negative effect on mRNA encapsulation, where larger particles (lower TFR) had lower mRNA encapsulation efficiencies (**Figure 4G**). The size controlled LNPs were then dosed in THP-1s based on encapsulated mRNA to elucidate differences in their ability to transfect macrophages. LNPs formulated at the highest TFR, thus the smallest LNPs generated, greatly outperformed the LNPs formulated at the lowest TFR in terms of macrophage mRNA expression, with nearly a 2-fold difference for all excipient formulations tested (**Figure 4G**). When interpreted alone, the results of the sizing characterization and subsequent screening suggest that smaller LNPs have a greater ability to deliver mRNA to macrophages. However, the additional trend with mRNA encapsulation efficiency suggests that size alone is not responsible for disparities across the different TFRs, and there may be additional LNP characteristics not adequately captured but may be indicative of LNP quality. Compared to higher TFRs, lower TFRs lead to slower mixing between the mRNA and lipid components, potentially leading to differences in the intermolecular packing of the lipids and mRNA within the particle itself, an aspect of LNP characterization that is gaining appreciation with more advanced characterization techniques.^88,89^ Thus, for two LNPs consisting of identical lipids and lipid compositions but formulated under different mixing conditions, the subsequent LNPs could have different internal organizations that cause significant deviations in performance. Further, the inability to achieve these packings may result in less efficient overall mRNA encapsulation, as demonstrated here. This could be a result of mRNA being a large and rigid molecule, thus it requires specific flow conditions for optimal packing within LNPs. Although these phenomena should be explored further, for the purposes of LNP optimization, the TFR for formulation was set to 2.4 mL/min. Thus, the optimized formulation moving forward was the B15 formulation, formulated at a 10:1 ionizable lipid:mRNA (wt/wt) under a TFR mixing condition of 2.4 mL/min.

Lastly, given the results of our DoE and weight ratio-based optimization of LNPs, it became clear that there were potentially two routes toward improving the ability of LNPs to deliver mRNA to macrophages: excipient optimization and weight ratio optimization. Thus, we took optimal conditions for both parameters (17.5:1 ionizable lipid:mRNA and the B15 excipient molar ratios) and applied them to three other LNPs from the previous ionizable lipid library screen which were either structurally similar to the C16-C lipid, or entirely different but had bioactivity in THP-1s. We found that the molar ratios used within the B15 formulation improved delivery for ionizable lipids containing the same polyamine core, but weight ratio-based optimization was more effective for different polyamine cores. Importantly, LNPs that had little to no activity in the initial screen (C14-B), had no improvement in mRNA delivery by either optimization route, further highlighting the role of the ionizable lipid, either by abundance or by identity, as a primary determinant for mRNA delivery to macrophages (**Supplementary** Figure 3).

### Extracellular pathways associated with LNP uptake and mRNA delivery to macrophages

Macrophages are highly specialized for the uptake of foreign extracellular materials and various serum components such as complement proteins, cholesterol, and lipoproteins.^90^ Internalization can occur through a variety of pathways including those that rely on cognate receptor-ligand interactions and via non-specific processes such as phagocytosis.^90–93^ It is well known that LNPs readily bind serum proteins including complement proteins and lipoproteins, and also contain cholesterol as a structural component.^90,94–97^ Thus, understanding the extracellular LNP-macrophage interaction can give insight into how macrophages perceive LNPs, a fundamental aspect of macrophage-LNP interactions. We explored the role of scavenger receptors (MARCO, CD204, CD36, and CD68) which aid in the clearance of pathogens and extracellular debris and have strong binding affinities for cholesterol and lipoproteins^98–100^, scavenger-independent lipoprotein receptors (LDLR), and complement receptors (CD11b, CD18, and MAC-1) which bind sequestered complement proteins^85,101,102^. To explore this, we utilized an antibody-mediated receptor blockade against each of these receptors under serum free (0% v/v), low serum (1% v/v), and full serum (10% v/v) conditions.

The antibody-mediated receptor blockade yielded several insights into the extracellular interactions that govern LNP mediated mRNA delivery to macrophages. Surprisingly, inhibition of the scavenger receptors, which bind lipoproteins and cholesterol, such as apolipoprotein E (ApoE), only led to modest decreases in mRNA delivery (< 50% knockdown) (**Figure 5A**). Of the lipoprotein-binding receptors tested, only LDLR and CD204 had serum-dependent effects on antibody-mediated blockade. For LDLR, increases in serum concentration led to increases in the inhibitory activity of LDLR blockade, and thus decreases in LNP bioactivity. For CD204, there was an inverse relationship, where increases in serum concentration led to decreases in the inhibitory activity of the receptor blockade. LDLR binds specifically to ApoE and, to a lesser extent, apolipoprotein B (ApoB).^103,104^ Thus, as serum concentration increases, more ApoE will adsorb to the LNP and bias LNP-macrophage interactions towards the LDLR and subsequent inhibition of this pathway will be more potent. CD204 also binds ApoE, albeit to a lesser extent, but also binds a wide range of additional ligands, thus by decreasing abundance of the ligands adsorbed to the surface to the LNP, the receptor blockade will be more effective at lower serum concentrations. Further, scavenger receptors of the same class can compensate for one another, that is, when one is occupied or inactive, other isoforms will be more active to maintain their overall function.^100,105^ Complement receptor inhibition, however, was highly serum dependent, as the inhibitory effect was almost entirely ameliorated in serum free media. These results indicate that complement receptors and lipoprotein both contribute to LNP-mediated mRNA delivery to macrophages.

**Figure 5:**
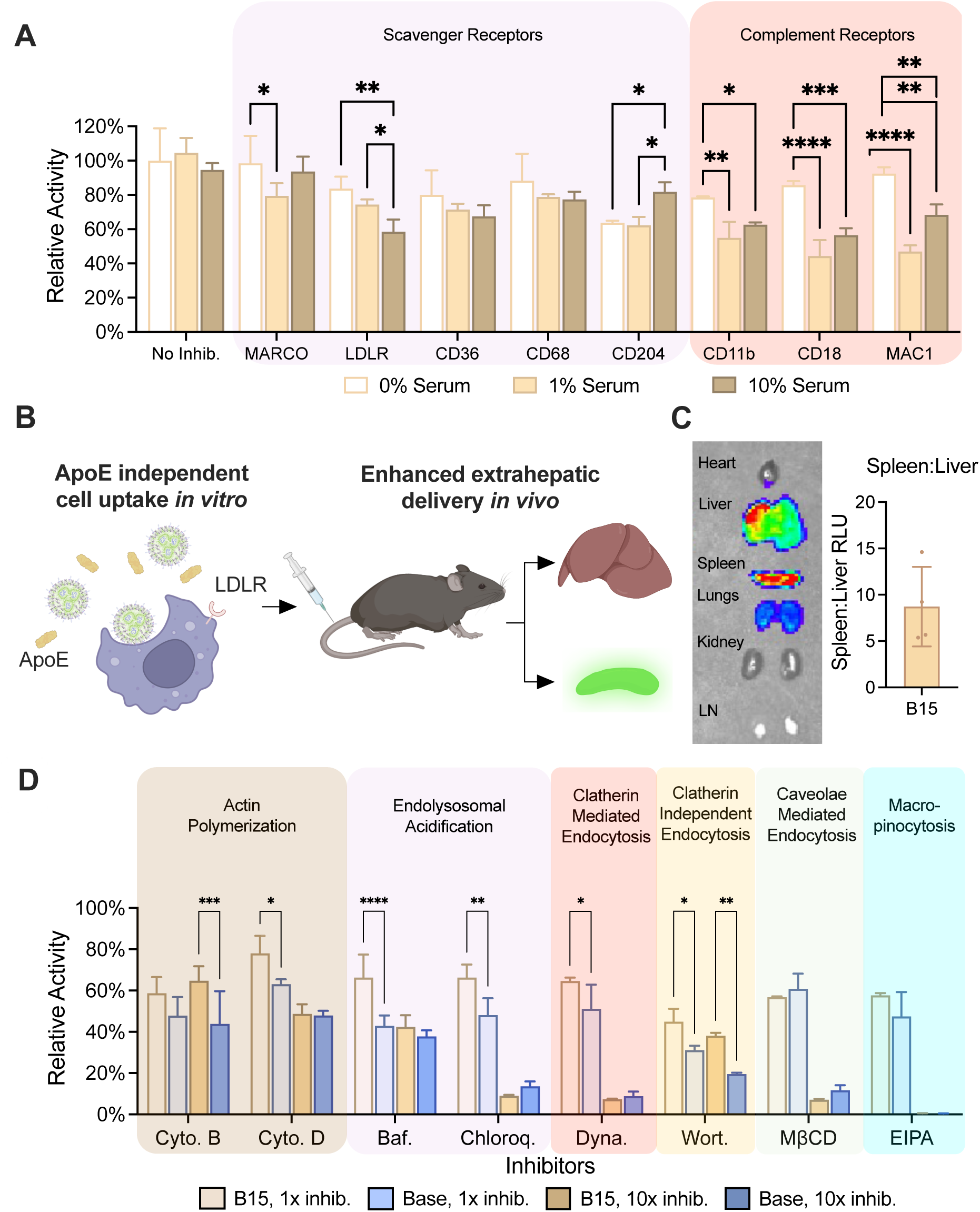
Optimized bioinspired mRNA-LNPs primarily deliver to macrophages through ApoE-independent macropinocytosis. (A) PMA-differentiated macrophages were incubated with antibodies for 2 hours at the indicated serum concentrations and subsequently treated with the optimized B15 formulation at a dose of 250ng/ 50k cells. Luminescence was measured 24 hours later and normalized to cells treated in antibody-free media. Data was analyzed using a 2-way ANOVA with Holm-Sidak correction for multiple comparisons with *n* = 4 biological replicates. (B-C) Mice were injected intravenously with the optimized B15 formulation encapsulating luciferase mRNA at a dose of 5 μg per mouse and luminescence signal for each organ was measured. Luminescence signal for each organ was first normalized to background before being normalized to liver signal in *n* = 4 mice. (D) PMA-differentiated macrophages were incubated with small molecule inhibitors of various endocytic pathways at a 1x and 10x doses for 2 hours and then treated with either the base formulation or the optimized B15 formulation at a dose of 250 ng/ 50k cells. Luminescence was measured 24 hours later and normalized to a parallel viability assay to account for inhibitor toxicity, before being normalized to uninhibited cells. Data was analyzed using a 2-way ANOVA with Holm-Sidak correction for multiple comparisons with *n* = 4 biological replicates.

To further confirm that LNP uptake is primarily ApoE independent, the optimized B15 LNP was used to encapsulate luciferase mRNA and administered intravenously to mice. LNPs were found to have significant tropism to the spleen, as evidenced by the high spleen:liver luminescence, contrary to the liver-dominant tropism observed with other strong ApoE-binding LNPs such as MC3 or C12-200 (**Figure 5B-C**).^106,107^ Of note, there was no singular inhibited receptor-ligand interaction that completely blocked LNP uptake, suggesting that that no specific adsorbed serum protein mediates LNP uptake into macrophages, and that LNP uptake might be driven by mechanisms largely independent of serum protein interactions.

### Small molecule inhibition of endocytic and phagocytic pathways associated with LNP uptake and mRNA delivery

Phagocytosis has long been considered a critical barrier for the intracellular delivery of exogenous cargos to macrophages, as the phagocytosis pathway is highly specialized for the degradation of uptake foreign materials. Although endocytosis is considered distinct from phagocytosis, they share similar intracellular trafficking pathways.^108–110^ That is, the transition from endosome to lysosome shares similar biochemical cues as the transition from phagosome to lysosome, namely in the acidification of these subcellular compartments, which occurs rapidly in macrophages. Notably, the ionizable lipid component of LNPs takes on positive charge at acidic pHs relevant to these pathways, making it an ideal carrier for RNA delivery to macrophages, as they might be able to escape phago-lysosomal trafficking in the same way that they escape endo-lysosomal trafficking. As such, we sought to elucidate some of the underlying mechanisms governing LNP trafficking within macrophages using selective inhibitors of various endocytic and phagocytic pathways. We constructed an inhibitor panel consisting of 8 small molecules that could inhibit the processes underpinning phagocytosis (cytochalasins B and D)^84,111–114^or inhibit the acidification of subcellular compartments (bafilomycin A and chloroquine).^115–117^ Further we included inhibitors of other endocytic pathways such as clatherin mediated endocytosis (Dynasore)^118,119^, clatherin independent endocytosis (wortmannin)^120,121^, caveolae mediated endocytosis (MβCD)^84,122,123^, and macropinocytosis (EIPA).^124,125^

The small molecule inhibitor assay first demonstrated that there is no singular pathway governing LNP-mediated mRNA delivery; several pathways contribute to varying degrees **(Figure 5D**). Further, the optimized B15 formulation maintained higher activity across several inhibitors and concentrations, confirming its increased potency compared to the base formulation. Notably, actin-polymerization, the process most discretely associated with phagocytosis in this panel, had the least impact on mRNA delivery, as delivery was still relatively high at the 10x inhibitor dose (<60% knockdown). Other inhibitors of endocytosis more profoundly inhibited mRNA delivery at the 10x dose, suggesting that endocytosis is the primary driver for the LNP-mediated intracellular delivery of mRNA. Interestingly, of all the molecules tested, EIPA, a selective inhibitor of macropinocytosis, was the only inhibitor to completely knockdown mRNA delivery. Macropinocytosis is a non-specific form of endocytosis that is constitutively active in macrophages, indicating that non-specific, fluid-phase uptake plays an important role in LNP-mediated mRNA delivery to LNPs.

The combined results of the antibody and small molecule inhibitor screens suggest that LNPs enter macrophages through a combination of specific and non-specific processes. Interactions with serum proteins such as ApoE and complement proteins enable cognate receptor-ligand interactions and induce endocytosis. Non-specific uptake, predominantly through macropinocytosis, simultaneously provides a route of entry for LNPs into the cell, where biochemical queues such as endosomal acidification enable LNPs to ionize, facilitate endosomal escape, and release mRNA cargo into the cytosol. The potential implication of these results suggests a degree of tunability of these LNPs, particularly towards receptor-specific mechanisms of uptake at the surface of the cell and away from non-specific mechanisms which may not provide any level of specificity in more complex cell environments.

### Optimized B15-LNPs do not induce changes in macrophage phenotype

Recent studies on the efficacy of LNPs for immunoengineering, specifically vaccines, revealed an intrinsic immunogenicity associated with LNPs independent of the mRNA cargo.^71,126,127^ Additional studies have shown that LNPs, through their mechanisms of action that promote endosomal escape, can interact with intracellular inflammatory pathways, especially in cells of the mononuclear phagocyte system, such as macrophages.^128^ Here, we characterized phenotypic responses to LNPs in donor derived primary human macrophages. LNPs were incubated with primary human macrophages (n = 3 donors) to assess changes in phenotype using multiplex gene expression analysis (NanoString) using a custom-curated panel of more than over 200 genes related to pro-inflammatory response, alternative macrophage activation, extracellular matrix regulation, fibrosis, and angiogenesis. Hierarchical clustering of gene expression values revealed only minor differences between macrophages treated optimized B15 LNPs and untreated macrophages 24h after LNP treatment (**Figure 6A-B**). Moreover, only eight genes were expressed significantly differently between both groups, when using an adjusted p-value cutoff of 0.05, and no genes were significantly different when using an adjusted p-value cutoff of 0.05 (**Figure 6B**). In contrast to previous findings describing the inflammatory effect of LNPs, B15 LNPs did not promote a pro-inflammatory response but instead decreased the expression of the inflammatory related genes IRF1 and IL6 (**Figure 6C**) and increased the expression of CD273, AFFG1, and FOXO1 (**Figure 6D**), markers of alternative macrophage activation.^129^ Of studies that have characterized immune responses to LNP vehicles, many have examined cytokine responses directly *in vivo*, thus, much of the immunological crosstalk and complexity captured in those studies is not recapitulated here.^127^ However, in general, the B15 formulation was not found to induce significant proinflammatory responses in donor-derived macrophages.

**Figure 6:**
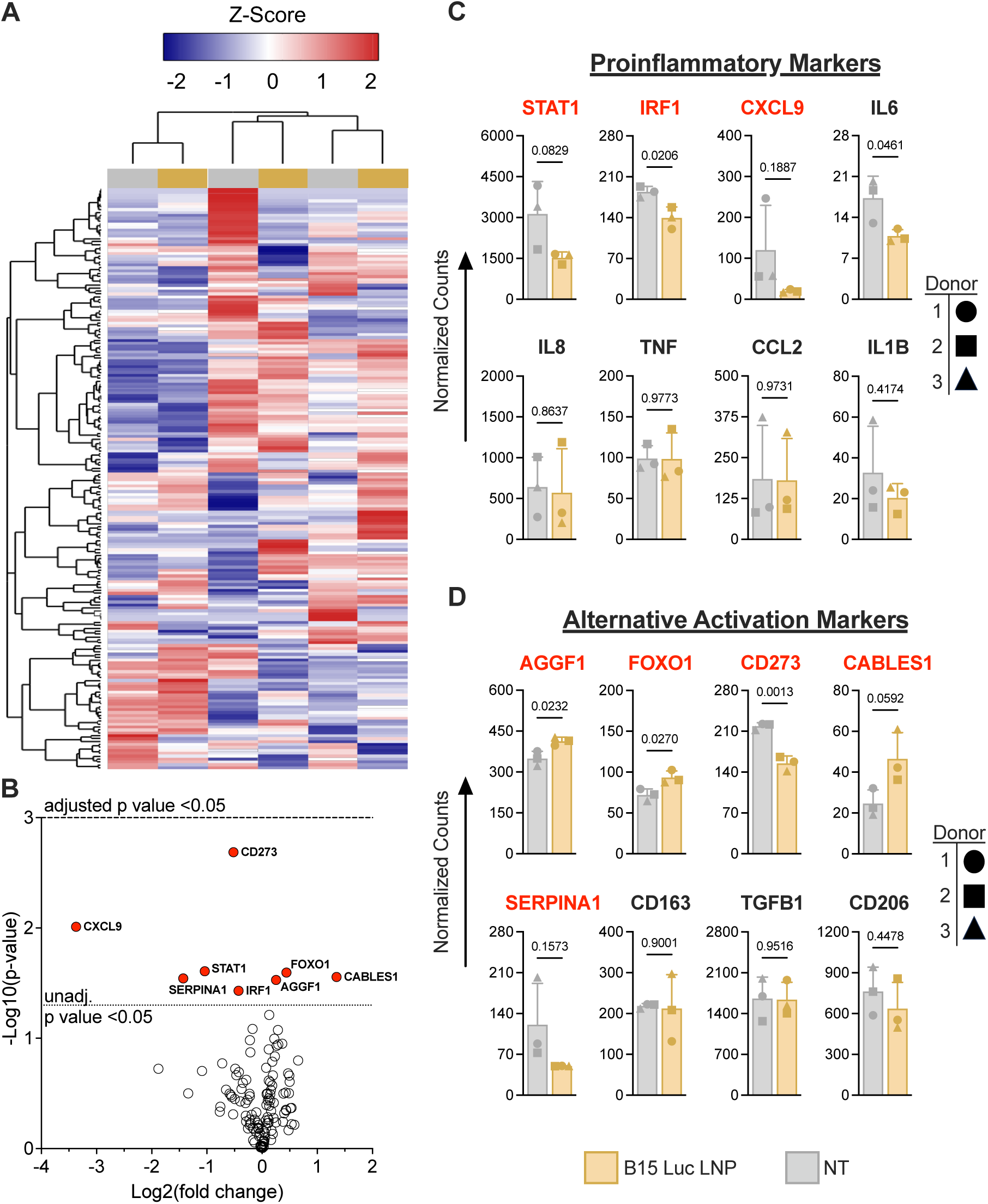
Optimized bioinspired mRNA-LNPs do not elicit a significant phenotypic response in donor-derived primary human macrophages. NanoString gene expression analysis of donor-derived primary human macrophages treated with the B15 luciferase mRNA-LNP at a dose of 500 ng/ 50k cells for 12 hours. (A) Hierarchical clustering of samples and genes in the heatmap represent Z-scores of all genes tested via a custom-made NanoString panel of either untreated (NT) or LNP-treated macrophages 24h after LNP treatment. (B) Volcano plot showing differential expressed genes (DEG) using an unadjusted p-value cutoff of 0.05 (shown as red dots and labeled); there were no DEGs when using an adjusted p-value cutoff of 0.05 (adjusted using a Benjamini-Yekutieli false discovery analysis). (C-D) Normalized counts of DEGs identified in (B) (red font) and additional representative (black font) (C) pro-inflammatory (IL6, IL8, TNF, IL1B, CCL2) and (D) alternative activation related markers (CD206, CD163, TGFB1). Data was analyzed using student t-test with *n* = 3 independent donors.

### Validation of optimized B15 LNP in MCSF- and GMCSF-*ex vivo* models of primary human macrophages

After exploring optimization routes and mechanisms of action for the C16-C LNP, the B15 formulation (10:1 weight ratio, 2.4 TFR) was chosen and studied in patient derived primary human macrophages to assess the translatability of this platform. Primary human monocytes were isolated from peripheral blood from the Human Immunology Core and the Hospital of University of Pennsylvania and cultured in media supplemented with granulocyte macrophage colony stimulating factor (GMCSF) or macrophage colony stimulating factor (MCSF) (**Figure 7A**). GMCSF- and MCSF-derived are the two most common macrophage models and are used to approximate inflammatory and unactivated macrophages respectively. Thus, both models were studied to assess if this LNP platform could be applied to both cell populations and if there were any quantifiably different patterns of LNP transfection and mRNA delivery across these models.

**Figure 7:**
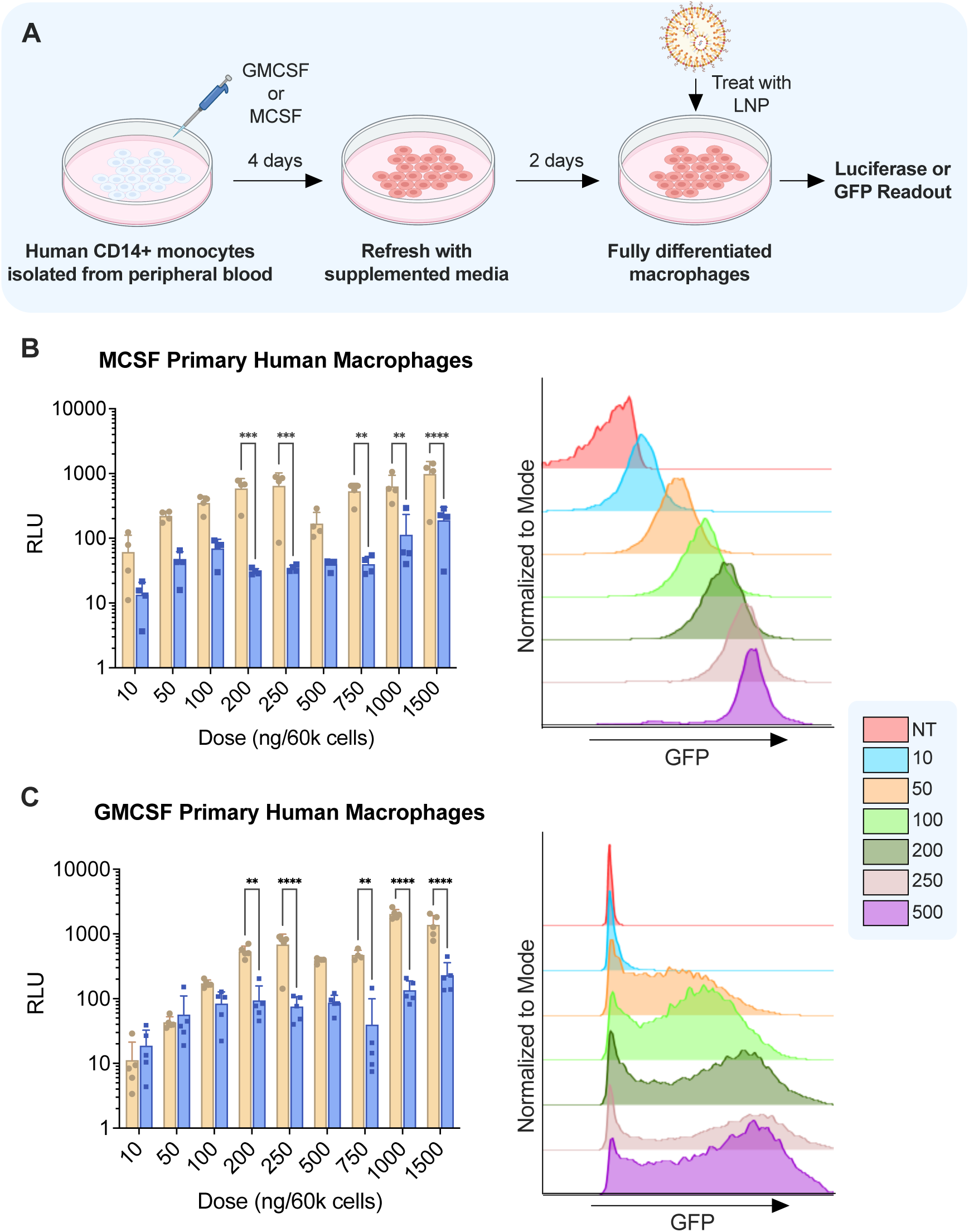
MCSF and GMCSF derived macrophages have differential responses to transfection with mRNA-LNPs. (A) Donor-derived CD14+ monocytes were differentiated into macrophages over the course of a 6-day *ex vivo* culture in MCSF or GMCSF supplemented media (20 ng/mL). MCSF-(B) or GMCSF-(C) primary macrophages were treated with luciferase mRNA LNPs (left) or GFP mRNA LNPs (right) at the indicated dose. Luminescence was measured 24 hours later. Data were normalized to untreated cells and analyzed using a 2-way ANOVA with Holm-Sidak correction for multiple comparisons with *n* = 4 independent donors. For GFP experiments, cells were harvested 24 hours after treatment with LNPs via gentle scraping and analyzed using flow cytometry with *n* = 4 independent donors and a single representative donor shown. . * p < .05, ** p < .01, *** p < .005, **** p < .001

The B15 and base formulations were compared using luciferase mRNA, where the B15 LNP was found to outperform the base formulation by nearly an order of magnitude in both models, confirming the findings of the DoE performed in THP-1s (**Figure 7B-C**). Next, the B15 LNP encapsulating GFP mRNA was studied in GMCSF and MCSF derived macrophages using flow cytometry to elucidate any differences in transfection efficiency. For GMCSF macrophages, GFP expression was heterogenous within a sample, as there was clear separation into GFP+ and GFP-cell populations, with a range of GFP-expressing cells in between (**Figure 7C**). In MCSF macrophages, nearly all cells within the sample were transfected at high doses. Interestingly, the GFP transfection profile of the GMCSF macrophages mirrored that of AdV transfection, which uses cognate receptor-ligand interactions for AdV cell entry and subsequent transgene expression (**Figure 7B-C**).^35^ The observed variability of GMCSF macrophages by both LNP and AdV suggests that GMCSF-derived macrophages are a more heterogenous population compared to MCSF-derived macrophages, and that the induced proinflammatory-like phenotype might interfere with exogenous mRNA expression, which has been previously shown.^130^ However, the B15 optimized LNP can be applied to either model, as it was able to induce robust mRNA expression resulting in significant MFI shifts in both GMCSF and MCSF macrophages.

### Oxidized B15 LNP can engineer functional HER2-CAR-Ms *ex vivo*

After confirming that the B15 formulation could successfully transfect primary human macrophages, we examined whether it could be used to generate functional primary human CAR macrophages (CAR-M). Clinically, GMCSF-macrophages are used in CAR-M therapies and were utilized here to highlight the clinical utility of this LNP platform. The B15 and Base formulations were used to encapsulate mRNA encoding a HER2-targeted CAR, similar in structure to one previously reported.^35^ After 24 hours, macrophages were harvested and HER2 CAR expression was measured using flow cytometry (**Figure 8A**). Macrophages were mixed with HER2+Luc+ SKOV3 human ovarian cancer cells at various effector:target (E:T, CAR+ Macrophage:SKOV3) ratios in a co-culture killing assay. After 48 hours, luciferase signal was quantified using a luciferase assay and SKOV3 killing was calculated by normalizing to SKOV3 cells cultured alone. Primary macrophages treated with B15 LNPs had a higher CAR positivity rate than macrophages treated with the base formulation (∼18% v. ∼3%) (**Figure 8B**). However, increases in transfection efficiency did not translate to statistically significant differences in the killing efficiencies between macrophages treated with the B15 or Base LNP formulation, which is due to the number of CAR+ macrophages added to the culture being the same for both groups. Although there was non-specific killing induced by untreated macrophages alone, both LNP formulations were able to achieve robust and dose-dependent tumor cell killing (**Figure 8C**). Although the CAR expression is lower than expected based on flow cytometry assays with GFP-encoding mRNA, it is likely that expression can be improved by further optimization of the CAR construct and the mRNA sequence. When treated with B15 LNPs encapsulating mRNA encoding a previously optimized CD19-CAR, CAR positivity increased to ∼66% compared to the 18% rate with the unoptimized HER2-CAR (**Supplementary** Figure 4). In sum, the improved transfection efficiency highlights the successful optimization for the C16-C particle. The improvement in efficiency is essential towards facilitating the translatability of mRNA-LNPs for CAR-M therapy.

**Figure 8:**
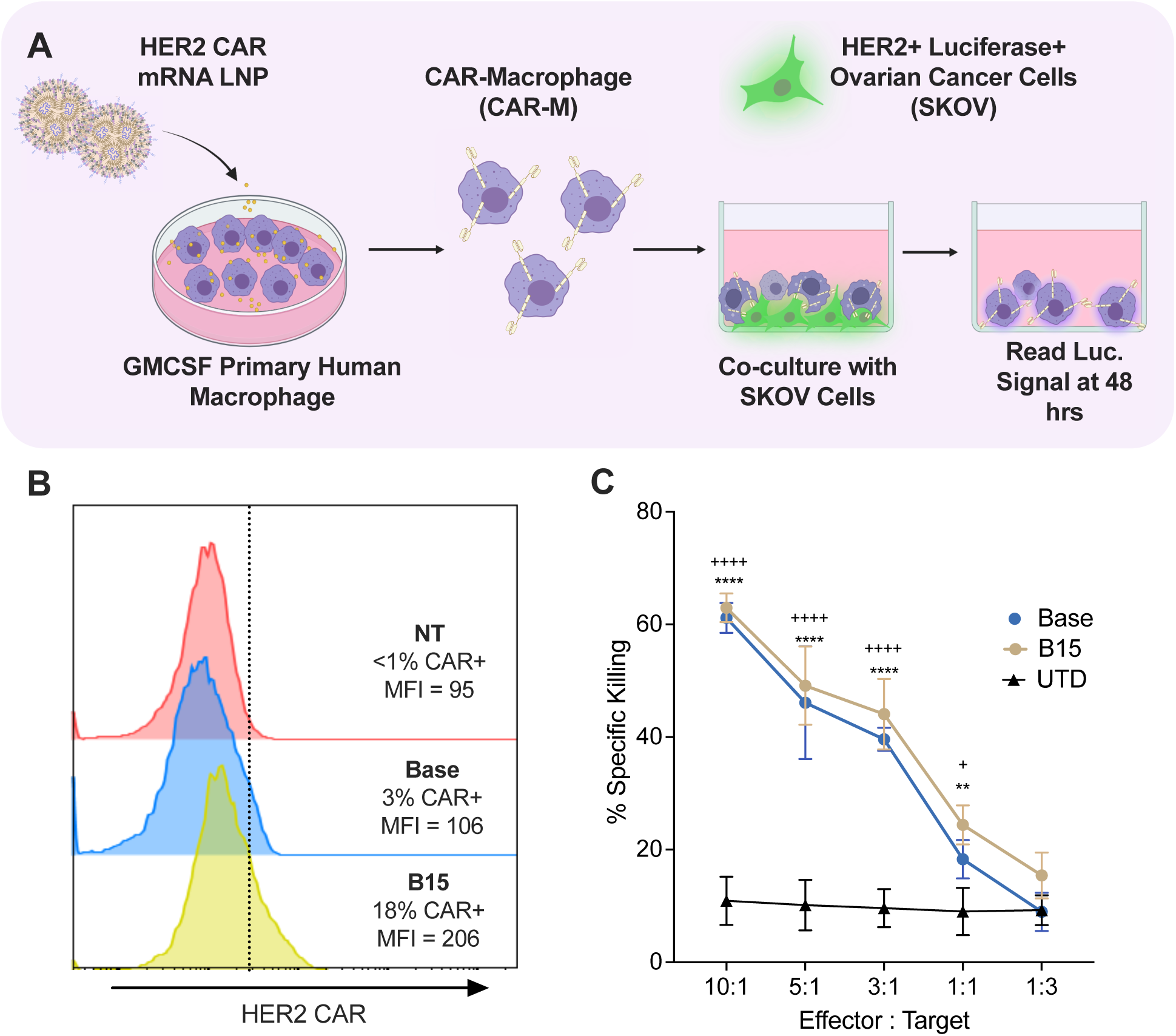
Oxidized B15 mRNA LNP engineers functional HER2-CAR macrophages *ex vivo*. (A) Experimental overview of *ex vivo* HER2 CAR macrophage-SKOV3 co-culture killing assay. (B) Donor-derived GMCSF macrophages were treated with base or B15 mRNA LNPs encapsulating HER2-CAR. CAR expression was measured after 24 hours using flow cytometry with *n* = 3 independent donors with a single representative donor shown. (C) HER2-CAR+ macrophages were incubated with HER2+ luciferase+ SKOV3 cells at various effector (CAR-M) to target (SKOV3) ratios with the total number of macrophages per well kept constant. After 48 hours, SKOV3 cell death was measured using a luciferase assay. Data was normalized to wells containing SKOV3 cells alone and compared using a 1-way ANOVA with *n* = 4 wells. B15 v. UTD:* p < .05, ** p < .01, *** p < .005, **** p < .001. Base v. UTD: + p < .05, ++ p < .01, +++ p < .005, ++++ p < .001.

## Conclusion

In sum, this work describes a new LNP platform for mRNA-based engineering of human CAR macrophages. Through an initial lipid screen, we identified the lipid C16-C capable of robust mRNA delivery to human macrophages *in vitro*, compared to a C12-200 gold standard. The excipient ratio of the C16-C LNP was then optimized using orthogonal DoE, demonstrating that higher ionizable lipid content was more favorable for mRNA delivery to macrophages, with minimal effects on toxicity, ultimately yielding an excipient optimized B15 formulation. Furthermore, the conditions of the microfluidic formulation of LNPs were studied, and it was found that smaller LNPs with higher mRNA encapsulation efficiencies were more potent than larger LNPs with lower mRNA encapsulation efficiencies. To provide insight into the mechanisms of LNP uptake into macrophages, small molecule and antibody-mediated inhibitor studies were performed, revealing these optimized LNPs deliver mRNA to macrophages through ApoE-independent macropinocytosis, which is further demonstrated by the potent extrahepatic tropism of the particle *in vivo*. To demonstrate the translatability of this platform we studied the potency of the optimized LNP in two common *ex vivo* models of primary human macrophages, GMCSF- and MCSF-derived macrophages, and found that although the optimized LNP was capable of potent mRNA, there were salient differences in the delivery profiles between the models, an important consideration for future studies. Lastly, we confirm that our optimized LNPs platform can engineer functional and potent human HER2-CARMs *ex vivo*, with efficiencies that can be enhanced with improved mRNA design. The flexibility of this platform enables the engineering of CAR-Ms against a broad range of CAR targets to treat a range of solid tumors and beyond. Furthermore, although we focus on engineering solid tumor-targeted CAR-Ms in this study, this platform can be used in other disease contexts where macrophages are heavily implicated, such as wound healing, atherosclerosis, and metabolic disorders.

## Materials and Methods

### Ionizable Lipid Synthesis

Epoxide-terminated alkyl chains were mixed with polyamine cores at a 7:1 molar ratio and allowed to react for 48 h at 80 °C in an excess of ethanol. The product was dried using a Rotavapor R-300 (Buchi) to remove excess solvent, before being resuspended at a concentration of concentration of 40 mg/mL in ethanol.

### Lipid Nanoparticle Formulation (LNP)

LNPs were formulated using microfluidic mixing of a lipid-containing ethanol phase and a mRNA-containing aqueous as previously described. For screening of the ionizable lipid library, the ethanol phase was prepared by combining ionizable lipid, DOPE (Avanti), cholesterol (Thermo), or a lipid anchored polyethylene glycol (C14-PEG_2000_) (Avanti) at a molar ratio of 35:16:46.5:2.5, respectively. The aqueous phase was prepared by diluting the appropriate mRNA to a concentration of .075 mg/mL in 10 mM citrate buffer (pH = 4). Ethanol and aqueous phases were mixed at 1:3 ratio (vol/vol) using single channel microfluidic mixing device. LNPs were subsequently collected in a 20 kDa MWCO dialysis cassette (Thermo), dialyzed against 1x PBS for 2 h, and sterile filtered using a .22 μm syringe filter. LNPs were stored in solution at 4C.

### Lipid Nanoparticle Characterization

#### LNP size, mRNA concentration, and mRNA encapsulation efficiency

For hydrodynamic radius and polydispersity measurements, LNPs were diluted 1:100 in PBS and analyzed in triplicate using dynamic light scattering (DLS) on a Malvern Zetasizer. mRNA content in LNP suspensions was measured using a Nanoquant plate (Tecan) and reported as the ratio of the absorbance measurements at 220 and 280 nm using a plate reader (Tecan). To measure mRNA encapsulation efficiency, a Quant-it RiboGreen assay kit was used to according to manufacturer protocols.

#### pKa Measurements

LNP pKa was determined using a 2-*p*-toluidinonaphthalene-6-sulfonate (TNS)-based fluorescence assay, as previously described.^131^ Briefly, LNPs were diluted to a concentration of 20 ng uL^-1^ in PBS, and 2.5 μL of this stock was diluted into well containing 100 μL of phosphate buffer at pH ranging from 3.0 to 12.0. 5 μL of TNS reagent was added to each well, incubated for 10 min, and the fluorescence signal of each well was measured at an excitation and emission of 325 nm and 450 nm respectively. Fluorescence values were normalized to the highest value and fitted to a sigmoidal function and pKa was determined to be pH at which the inflection point occurred.

### Orthogonal Design of Experiments

*Library Generation*: Orthogonal design of experiment LNP libraries were generated using JMP software. For library A, molar ratio ranges for ionizable lipid and each excipient was based on the screening formulation and each excipient molar ratio was changed at 4 level. For library B, molar ratio ranges were higher and based on the A12 formulation, and each excipient molar ratio, aside from PEG-Lipid, was changed at 4 levels. Based on the trends from library A, the PEG-Lipid molar ratio was held constant at 2.5

### Cell Culture

#### THP-1 Cell Culture

THP-1 monocytes were cultured in suspension in RPMI supplemented with 10% Heat Inactivated FBS and 1% penicillin-streptomycin at a density of 2.5 x 10^5^–2.0x10^6^ cells/mL.

#### Primary Human Macrophage Cell Culture

Primary human monocytes (CD14+) were collected from the peripheral blood of healthy human donor patients through the UPenn Human Immunology Core. On day 0 *ex vivo* culture was initiated and monocytes were plated directly into 96-well plates for luciferase and toxicity assay or 24-well plates for flow cytometry and killing assays. Monocytes were cultured in Glutamax RPMI supplemented with 10% Heat Inactivated FBS, 1% penicillin-streptomycin and GMCSF (20 ng mL^-1^) (Peprotech, 300-03) or MCSF (20 ng mL ^-1^) (Peprotech, 300-25). Differentiation into macrophages was monitored through the adherence of cells. On day 3, supernatant, containing undifferentiated monocytes, were collected, spun down, resuspended in GMCSF- or MCSF-supplemented culture media and added back to the wells. mRNA-LNPs were added to cells on day 6.

#### mRNA Delivery to THP-1 Macrophages

For luciferase assays, cells were spun down and resuspended to a concentration of 5.0x10^5^ in culture media further supplemented with 10 ng/mL of phorbol 12-myristate 13-acetate (PMA). 100 μL of the suspension was added to each well of a 96-well plate for a final cell concentration of 5x10^4^ cells per well. Cells were differentiated for 48 hours before cells were refreshed with PMA-free media to remove PMA and undifferentiated cells. After 1 hr, LNPs encapsulating luciferase-encoding mRNA were added at the indicated concentrations. After 24 h of incubation with the LNPs, luciferase signal was measured using a Luciferase Assay (Promega) according to manufacturer protocols. Briefly, media was aspirated off and 50 μL of lysis buffer was added, incubated for 10 min, before 100 μL of luciferase assay substrate was added. The assay was incubated in the dark at room temperature for 10 min before being read on Infinite M plex plate reader (Tecan, Morrisville, NC). Background luminescence signal was calculated from wells without cells but containing lysis buffer and luciferase assay reagent and subtracted from cell-containing wells. For analysis, luminescence signal for each group was first normalized to untreated cells and then normalized to the luminescence signal of cells treated with C12-200 LNPs.

For toxicity assays, cells were plated using the same protocol and treated with either C16-C LNPs, Lipofectamine 2000 transfection reagent, or C12-200 at dose of 500 ng of mRNA. Lipofectamine 2000 reagent was prepared according to manufacturer’s protocol. After 24 h, equal volume of CellTiterGlo (Promega) reagent was added and luminescence signal was measured using a plate reader. Background signal was calculated from cell-free wells containing media and assay reagents and was subtracted. Luminescence signal was normalized to untreated cells, with untreated cells considered to have 100% viability.

#### mRNA Delivery to GMCSF and MCSF Primary Human Macrophages ex vivo

For luciferase and toxicity assays, cells were diluted to 5 x 10^5^ cells/mL in growth media supplemented with 10 ng/mL of GMCSF or MCSF and plated in a 96-well plate on day 0. Cells were refreshed with GMCSF- or MCSF-supplemented growth media on day 3 and treated with LNPs encapsulated luciferase-encoding mRNA on day 6 as described above. After 24 h, luciferase or CellTiterGlo toxicities assays were performed according to manufacturer protocols previously described for THP-1s.

For flow cytometry studies, on day 0, cells were diluted up to 2.5 x 10^5^ cells/mL in GMCSF- or MCSF-supplemented media and plated at 2.5 x 10^5^ cells/well in a 24-well plate. Cells were refreshed with fully supplemented media on day 3 and treated with LNPs encapsulating GFP encoding mRNA on day 6 as described above. After 24 h cells were non-enzymatically lifted using a combination of CellStripper (Corning) and gentle scraping, spun down, and resuspended in PBS supplemented with .1% bovine serum albumin (BSA). Cells were analyzed on a BD LSR II Flow Cytometer using FACS Diva software, and the GFP signal was analyzed for 10,000 single cell events using FlowJo. To determine HER2-CAR expression for primary human macrophages treated with HER2-CAR mRNA LNPs, cells were plated and lifted as described above. After resuspension in PBS + .1% BSA, cells were stained with HER2 protein fragment tagged with a poly-histidine tail (Sino Biological). After 30 min, cells were rinsed with PBS + 1% BSA, and incubated with Human TruStain FTX Fc-blocking reagent for 10 min before a fluorescently labeled with APC anti-His secondary antibody was added and the entire mixture was incubated for 20 min. Fully stained cells were rinsed before being resuspended in PBS + .1% BSA and analyzed on BD LSR II Flow Cytometer using FACS Diva software. HER2-CAR signal was analyzed for 10,000 single cell events using FlowJo.

#### Flow rate

For mRNA LNPs formulated at various flow rates, the same formulation was mixed at different rates, but the 1:3 EtOH:Aq (v/v) ratio was held constant. Flow rates were changed on the Harvard Apparatus 3000 Syringe. The total flow rates (TFRs) were equivalent to the sum of the individual input flow rates (FR1+FR2).

| Aqueous Phase<br>(mL min<sup>-1</sup>) | Lipid Phase<br>(mL min<sup>-1</sup>) | Total Flow Rate<br>(mL min<sup>-1</sup>) | Flow Rate Ratio<br>(Aqueous: Lipid) |
| --- | --- | --- | --- |
| 0.18 | 0.06 | 0.24 | 3:1 |
| 0.45 | 0.15 | 0.60 | 3:1 |
| 0.90 | 0.30 | 1.20 | 3:1 |
| 1.80 | 0.60 | 2.40 | 3:1 |

#### Antibody mediated receptor blockade

For extracellular inhibition of surface receptor studies, THP-1s monocytes were plated and PMA-differentiated into macrophages in a 96-well plate for 48 h as previously described. 30 min before treatment with luciferase mRNA LNPs, one test, as specified by the manufacturer, of LDLR (Invitrogen, Cat: MA532075), CD18 (Invitrogen, Cat: 371300), CD11b (Invitrogen, Cat: MA180091), CD36 (Invitrogen, Cat: PA1-16813), CD68 (Invitrogen, Cat: 14-06888-82), MARCO (Invitrogen, Cat: PA5-64134), CD204 (Invitrogen, Cat: 14-9054-82) or MAC-1 (CD11b + CD18, 1:1) antibodies were added to THP-1 macrophages to block receptor interactions. Following the 30 min preincubation with antibodies, B15 LNPs formulated at a 2.4 mL min^-1^ TFR and 10:1 ionizable lipid:mRNA (wt:wt) encapsulating luciferase mRNA were added to each well at an mRNA dose of 200 ng well^-1^. After 24 h incubation, a luciferase assay was performed according to manufacturer’s protocols. In parallel, a duplicate plate was treated under the exact procedure and toxicity was measured using a CellTitrGlo kit according to manufacturer’s protocols. Luminescence signal was normalized to toxicity values to account for differences between groups occurring as a result of cell death.

#### Small molecule endocytosis and phagocytosis inhibitor assays

For intracellular inhibition of endocytosis and phagocytosis studies, THP-1 monocytes were plated and PMA-differentiated into macrophages in a 96-well plate for 48 h as previously described. The inhibitors: 5-(N-ethyl-n-isopropyl)-amiloride (EIPA, 10 μM, 100 μM), methyl-β-cyclodextrin (MβCO, 1 mM, 10 mM), wortmannin (1 μM, 10 μM), cytochalasin D (1 μM, 10 μM), cytochalasin B (1 μM, 10 μM), bafilomycin A1 (.2 nM, 2 nM), and chloroquine (20 μM, 200 μM) were dissolved in DMSO to make 100x concentrated stock solutions. 2 h prior to treatment with mRNA LNPs, each inhibitor was diluted 100x into macrophage-containing wells in a 96-well plate (1 μL). Additionally, a set of replicates of cells were treated with 1 μL of inhibitor-free DMSO to account for differences in LNP-mediated mRNA delivery because of enhanced cell permeability due to the presence of DMSO. After 2 h of incubation, cells were treated with the Base or the B15 LNP formulation encapsulating luciferase mRNA at an mRNA dose of 200 ng well^-1^. After 24 h of treatment with LNPs, a luciferase assay was performed according to manufacturer protocols. Luminescence signal was normalized to toxicity values to account for differences in luminescence between groups occurring as a result of cell death.

| Inhibitor | Final Well Concentration<br>(1x Dose) | Final Well Concentration<br>(10x Dose) |
| --- | --- | --- |
| EIPA | 10 μM | 100 μM |
| MβCO | 1 mM | 10 mM |
| Wortmannin | 1 μM | 10 μM |
| Cytochalasin D | 1 μM | 10 μM |
| Cytochalasin B | 1 μM | 10 μM |
| Bafilomycin A1 | .2 nM | 2 nM |
| Chloroquine | 20 μM | 200 μM |

#### NanoString Analysis

Primary human monocytes were isolated from peripheral blood of one donor via gradient centrifugation and cultured in primary media (RPMI-1640 (Thermo Fisher Scientific, Waltham, USA)+ 10% human serum + 1% penicillin/streptomycin) supplemented with macrophage colony stimulating factor (MCSF) (Peprotech, Cranbury, USA) for four days to induce macrophage differentiation, with a media change on day three. On day four, media was refreshed and one group received additional supplementation of lipopolysaccharide and interferon-gamma (inflammatory media) to induce pro-inflammatory macrophage polarization. On day five, unactivated and pro-inflammatory macrophages were co-cultured in triplicate with blank LNPs for 12 hours to facilitate phagocytosis, at a dose of 500ng per 50,000 cells. Some macrophages from each group were not co-cultured with LNPs to act as untreated controls. After 2 or 12 hours, excess LNPs were washed off. 24 hours later, cells were lysed and RNA was extracted using the RNAqueous-Micro Total RNA Isolation Kit (Thermo Fisher, Waltham, USA). 100ng of RNA from each sample was hybridized with a custom NanoString Codeset of over 200 genes (NanoString Technologies, Seattle, USA), including markers for pro-inflammatory, alternative-activated macrophage phenotype. Gene counts were measured on the NanoString nCounter and normalized to internal controls. T-tests were used to compare the gene expression of either group against its baseline control. Volcano plot analysis was performed in nSolver software with Benjamini-Yekutieli false discovery analysis. Hierarchical clustering was performed in R based on z-scores of normalized gene counts per gene.

### Animal Experiments

#### In vivo biodistribution imaging

6-8 week old C57BL/6 mice were injected via the tail vein at a dose of 5 ug luciferase mRNA per mouse with the optimized B15 formulation. 6 hr later, mice were injected intraperitoneally with 200 μL luciferin salt solution (15 mg/mL). After 10 min, mice were sacrificed and the heart, lungs, liver, spleen, kidneys, and inguinal lymph nodes were harvested and imaged using an IVIS. For quantification, individual ROIs were drawn around each organ and signal was normalized to the overall background signal of the image.

### HER2 CAR-Macrophage *ex vivo* Killing Assay

CAR macrophages were added to wells containing adherent HER2+ Luciferase+ SKOV3 human ovarian cancer cells at increasing effector (macrophage) to target (SKOV3) ratios. The total number SKOV3 cells per well and total number of macrophages per E:T ratio was kept constant across treatment groups for direct comparison. SKOV3 cell killing was determined by comparing luciferase signal in wells receiving macrophages to wells containing SKOV3 cells alone, and killing was compared between HER2+CARMs and untreated, CAR-macrophages (UTD).

