## Supplementary Data for "Bioinspired Oxidized mRNA Lipid Nanoparticles for *Ex Vivo* Engineering of Chimeric Antigen Receptor Macrophages Targeting Solid Tumors"

### Supplementary Data for Manuscript Titled: Bioinspired Oxidized mRNA Lipid Nanoparticles for *Ex Vivo* Chimeric Antigen Receptor Macrophages Targeting Solid Tumors

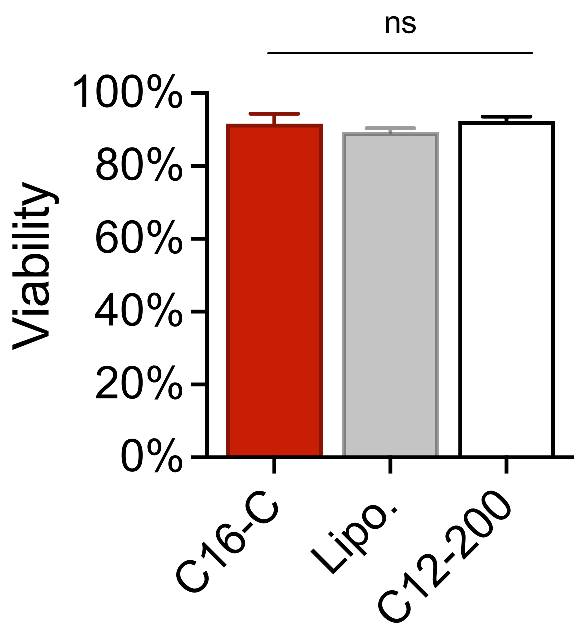

**Supplementary Figure 1**. Toxicity of the lead C16-C LNP was found to be not significantly different than toxicity associated with other gold standard mRNA delivery vehicles Lipofectamine and C12-200 LNP. Viability was measured using a CellTiter Glo assay in PMA-differentiated THP-1 macrophages after 24 hr incubation with the indicated treated. Data was normalized to untreated cells.

**Library A**

| Formulation | C16-C | DOPE | CHOL | PEG Lipid |
|---|---|---|---|---|
| 1 | 5 | 4 | 46 | 3 |
| 2 | 5 | 16 | 31 | 2 |
| 3 | 5 | 28 | 16 | 1.5 |
| NULL | 5 | 40 | 61 | 2.5 |
| 4 | 20 | 4 | 31 | 1.5 |
| 5 | 20 | 16 | 46 | 2.5 |
| 6 | 20 | 28 | 61 | 3 |
| 7 | 20 | 40 | 16 | 2 |
| 8 | 35 | 4 | 61 | 2 |
| 9 | 35 | 16 | 16 | 3 |
| 10 | 35 | 28 | 31 | 2.5 |
| 11 | 35 | 40 | 46 | 1.5 |
| 12 | 50 | 4 | 16 | 2.5 |
| 13 | 50 | 16 | 61 | 1.5 |
| 14 | 50 | 28 | 46 | 2 |
| 15 | 50 | 40 | 31 | 3 |

**Supplementary Table 2.** Molar ratios for the Library A formulations screened in the first round of excipient DoE optimization

**Library B**

| Formulation | C16-C | DOPE | CHOL | PEG |
|---|---|---|---|---|
| 1 | 40 | 4 | 16 | 2.5 |
| 2 | 40 | 8 | 12 | 2.5 |
| 3 | 40 | 12 | 8 | 2.5 |
| 4 | 40 | 16 | 20 | 2.5 |
| 5 | 50 | 4 | 12 | 2.5 |
| 6 | 50 | 8 | 16 | 2.5 |
| 7 | 50 | 12 | 20 | 2.5 |
| 8 | 50 | 16 | 8 | 2.5 |
| 9 | 60 | 4 | 20 | 2.5 |
| 10 | 60 | 8 | 8 | 2.5 |
| 11 | 60 | 12 | 12 | 2.5 |
| 12 | 60 | 16 | 16 | 2.5 |
| 13 | 70 | 4 | 8 | 2.5 |
| 14 | 70 | 8 | 20 | 2.5 |
| 15 | 70 | 12 | 16 | 2.5 |
| 16 | 70 | 16 | 12 | 2.5 |

**Supplementary Table 3.** Molar ratios for the Library B formulations screened in the second round of excipient DoE optimization

| Formulation | Hydrodynamic Radius (nm) | PDI | mRNA Concentration (ng/μL) |
|---|---|---|---|
| Base | 84.15 ± 3.25 | 0.222 | 39.40 |
| A1 | 123.83 ± 1.11 | 0.205 | 56.92 |
| A2 | 102.83 ± 2.92 | 0.153 | 43.84 |
| A3 | 92.40 ± 1.56 | 0.151 | 39.92 |
| A4 | 96.97 ± 3.18 | 0.151 | 34.92 |
| A5 | 94.30 ± 2.30 | 0.211 | 30.92 |
| A6 | 116.60 ± 2.95 | 0.273 | 43.56 |
| A7 | 93 ± 3.33 | 0.098 | 33.48 |
| A8 | 105.78 ± 2.68 | 0.176 | 35.76 |
| A9 | 110.83 ± 4.68 | 0.276 | 45.84 |
| A10 | 92.30 ± 1.81 | 0.191 | 33.44 |
| A11 | 89.43 ± 4.65 | 0.052 | 46.24 |
| A12 | 70.62 ± 2.46 | 0.170 | 49.28 |
| A13 | 78.11 ± 2.14 | 0.162 | 26.20 |
| A14 | 82.37 ± 3.74 | 0.078 | 29.20 |
| A15 | 71.72 ± 5.13 | 0.159 | 33.56 |
| B1 | 85.91 ± 2.13 | 0.199 | 40.52 |
| B2 | 89.30 ± 1.19 | 0.079 | 52.34 |
| B3 | 76.50 ± 4.14 | 0.284 | 33.08 |
| B4 | 70.12 ± 5.91 | 0.147 | 41.72 |
| B5 | 82.56 ± 4.30 | 0.233 | 38.60 |
| B6 | 83.30 ± 2.20 | 0.194 | 35.84 |
| B7 | 77.33 ± 3.59 | 0.225 | 36.64 |
| B8 | 72.68 ± 2.12 | 0.186 | 34.42 |
| B9 | 81.75 ± 4.15 | 0.178 | 40.48 |
| B10 | 83.40 ± 2.46 | 0.238 | 48.92 |
| B11 | 79.26 ± 6.20 | 0.189 | 36.92 |
| B12 | 85.50 ± 3.89 | 0.259 | 37.92 |
| B13 | 75.65 ± 1.44 | 0.149 | 30.32 |
| B14 | 71.92 ± 1.85 | 0.177 | 32.32 |
| B15 | 78.52 ± 2.28 | 0.129 | 31.64 |
| B16 | 87.72 ± 2.75 | 0.061 | 41.12 |

**Supplementary Table 4.** Physicochemical characterization of LNP formulations tested in DoE libraries A and B. Hydrodynamic radius and PDI were measured using dynamic light scattering and mRNA concentration was measured using a NanoQuant plate

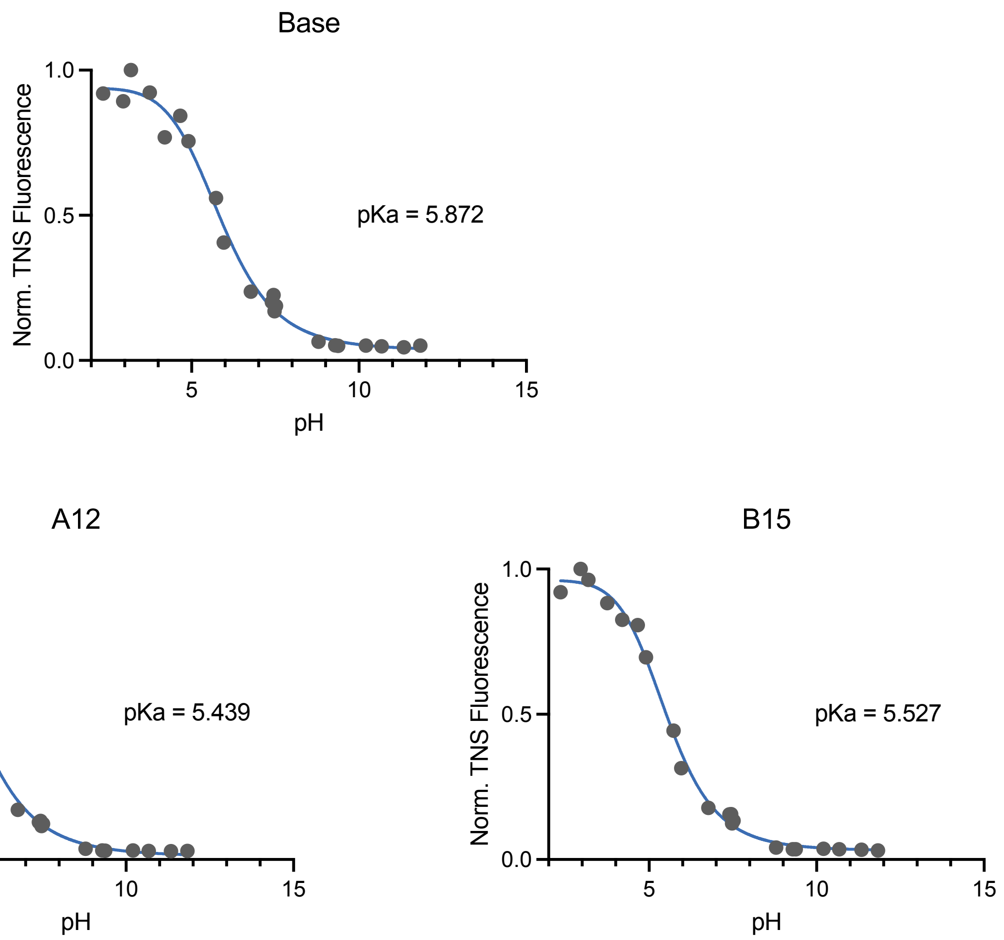

**Supplementary Figure 2.** TNS fluorescence curves used to calculate the pKa for the Base, A12, and B15 excipient optimized LNPs.

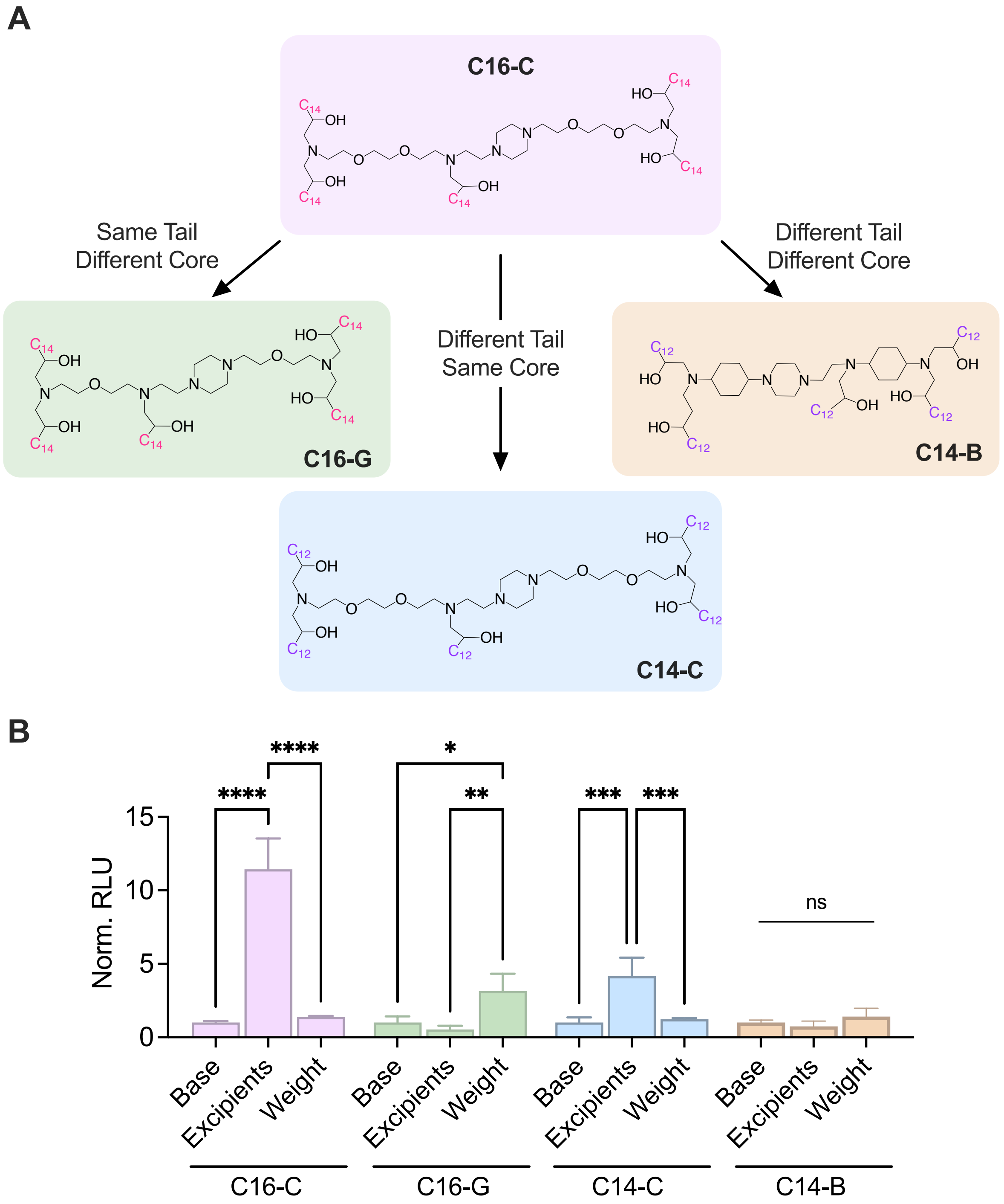

**Supplementary Figure 3: Optimization route of LNPs for mRNA delivery to macrophages is unique to each constitutive ionizable lipid.** (A) Three additional ionizable lipids from the 24 ionizable lipid library screened in Figure 2 were formulated using the base excipient ratios, B15 excipient ratios or the 17.5:1 ionizable lipid:mRNA (wt:wt) ratio, representative of an excipient-based approach or a weight ratio approach towards increasing the potency of mRNA LNP delivery to macrophages. (B) PMA-differentiated THP-1 macrophages were treated with luciferase mRNA LNPs at a dose of 250 ng/ 50k cells. Luminescence was measured 24 hours later. For each ionizable lipid, luminescence was normalized to the base group and compared using a 2-way ANOVA. n = 4 biological replicates. * $p < .05$, ** $p < .01$, *** $p < .005$, **** $p < .001$.

| | | | | | | | | |
|---|---|---|---|---|---|---|---|---|
| ABCG2 | CCL19 | CD68 | EGR2 | IL10RB | LXRbeta (NR1H2) | PFKFB1 | TIMP-1 | B-actin (ACTB) |
| ACOD1 | CCL2 | CD80 | ENO1 | IL12A | Lrg1 | PFKFB3 | TIMP-2 | CYC1 |
| ACTA2 | CCL20 | CD93 (C1qR1) | EPAS1 | IL12p40 (IL-12B) | MARCO | PKM2 | TIMP3 | EIF2B2 |
| AGGF1 | CCL22 (MDC) | CLEC10A | ERG | IL13RA1 | MCTP2 | PLOD2 | TIPE2 | GUSB (beta glucuronidase) |
| ALDOA | CCL24 | COL11A1 | ETS1 | IL13RA2 | MERTK | PPARD | TLR2 | HPRT1 (hypoxanthine guanine phosphoribosyl transferase) |
| ALOX15 | CCL26 | COL1A1 | FBP1 | IL17 | MEST | PPARg | TLR3 | TBP |
| ANG | CCL4 | COL3A1 | FLT1 (VEGFR1 ) | IL1B | MMP13 | PTGES2 | TLR4 | |
| APOBEC 3A. | CCL5 | COL5A1 | FN1 | IL1R | MMP2 | RAGE (AGER) | TLR7 | |
| ATG5 | CCL8 | CREB1 | FOXO1 | IL4 | MMP7 | RAMP1 | TNF | |
| Abca9 | CCN2 (CTGF) | CTNNB1 | FOXO3 | IL4Ralpha | MMP8 | RIPK3 | TNFAIP6 | |
| ApoE | CCR10 | CX3CR1 | FOXO4 | IL6 | MMP9 | S100a8 | TNFRII/C D120b | |
| Axl | CCR2 | CXCL10 | FST | IL8 | MRC1 (CD206) | SERPINA 1 | TNFRSF1 1A (RANK) | |
| BAX | CCR7 | CXCL11 | FSTI1 | IRF4 | MS4A6E | SH3PXD2 B | TNFRSF1 A | |
| BGN | CCR8 | CXCL12 | FYN | IRF5 | NAA15 | SOCS1 | TRAF6 | |
| BTG1 | CD11b | CXCL2 | GAL3 | ITGB1BP 1 | NFKB1 | SOCS3 | TYK2 | |
| Bcl2 | CD150/S LAM | CXCL3 | GATA3 | JAK2 | NNMT | SPHK1 | Trf | |
| C3AR1 | CD16 | CXCL9 | GLUL | JUN | NOD 1 | SPP1 | VCAN | |
| CABLES1 | CD163 | CXCR2 | HIF1A | Jag1 | NOD 2 | SREBF1 | VEGFA | |
| CACNA1 G | CD166 | CXCR4 | HLA-DRA | Jak1 | OLR1 | STAT3 | VEGFB | |
| CACNB4 | CD1C | CYR61 | HSPG2 (Perlecan ) | Jak3 | PDGFA | STAT6 | VEGFC | |
| CARKL | CD200R1 | Cox2 | IDO1 | KLF4 | PDGFB | Stat1 | VIM | |
| CCL1 | CD273 | DACT1 | IDO2 | LIGHT | PDGFC | TGFB1 | WNT5A | |
| CCL15 | CD274 | DCN | IGF1 | LIN7A | PDGFRA | TGFB3 | cd209 | |
| CCL17 | CD36/SR-B3 | DNASE1L 3 | IL-13 | LUM | PDGFRB | TGM2 | irf1 | |
| CCL18 | CD38 | EGFL7 | IL10 | LXRalpha (NRIH3) | PECAM1 | TIE1 | morc4 | |

**Supplementary Table 5:** List of genes studied in donor-derived primary human macrophages via Nanostring gene expression analysis in main text Figure 6.

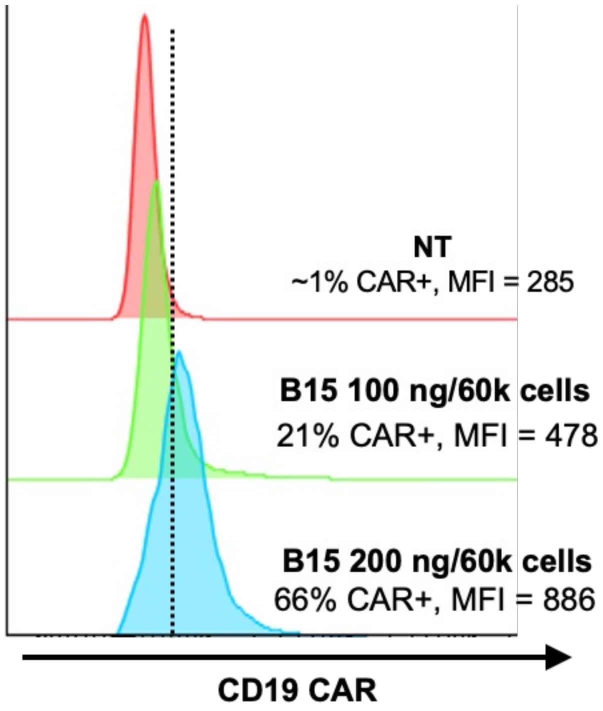

**Supplementary Figure 4: CD19-CAR mRNA is expressed more potently than HER2-CAR.** CD19-CAR was expressed by 66% of primary macrophages, compared to 18% expression of HER2-CAR at an equivalent dose (Main Text Figure 7B).
